# Receptor Allostery Promotes Context-Dependent Sonic Hedgehog Signaling During Embryonic Development

**DOI:** 10.1101/2025.01.28.635336

**Authors:** Shariq S. Ansari, Miriam E. Dillard, Mohamed Ghonim, Yan Zhang, Daniel P. Stewart, Robin Canac, Ivan P. Moskowitz, William C. Wright, Christina A. Daly, Shondra M. Pruett-Miller, Jeffrey Steinberg, Yong-Dong Wang, Taosheng Chen, Paul G. Thomas, James P. Bridges, Stacey K. Ogden

## Abstract

Sonic Hedgehog (SHH) signaling functions in temporal- and context-dependent manners to pattern diverse tissues during embryogenesis. The signal transducer Smoothened (SMO) is activated by sterols, oxysterols, and arachidonic acid (AA) through binding pockets in its extracellular cysteine-rich domain (CRD) and 7-transmembrane (7TM) bundle. *In vitro* analyses suggest SMO signaling is allosterically enhanced by combinatorial ligand binding to these pockets but *in vivo* evidence of SMO allostery is lacking. Herein, we map an AA binding pocket at the top of the 7TM bundle and show that its disruption attenuates SHH and sterol-stimulated SMO induction. A knockin mouse model of compromised AA binding reveals that homozygous mutant mice are cyanotic, exhibit high perinatal lethality, and show congenital heart disease. Surviving mutants demonstrate pulmonary maldevelopment and fail to thrive. Neurodevelopment is unaltered in these mice, suggesting that context-dependent allosteric regulation of SMO signaling allows for precise tuning of pathway activity during cardiopulmonary development.

## Introduction

Embryonic development relies on precise spatial and temporal coordination of cell behaviors to facilitate tissue morphogenesis. The Sonic Hedgehog (SHH) signaling pathway contributes to tissue organization by providing positional information and guiding cell fate decisions in signal-responding cells ^1^. Through this activity, SHH instructs body asymmetry and contributes to development of diverse organ systems including the nervous system, heart, lungs, digestive tract, and musculoskeletal system ^2–5^. Consistent with its role as a master regulator of tissue patterning and homeostasis, dysregulation of SHH signaling leads to developmental disorders such as holoprosencephaly (HPE), congenital heart disease (CHD), and lung hypoplasia, and aberrant signaling can drive tumor formation in the cerebellum and skin ^1,4–6^.

In most vertebrate cell types, the SHH signaling response is coordinated through the primary cilium (PC), a specialized sensory organelle that extends from the basal body and facilitates signaling by members of the G protein-coupled receptor (GPCR) superfamily ^7,8^. In the absence of ligand, the SHH receptor Patched (PTCH) localizes to primary cilia and depletes sterols from ciliary membrane ^9–11^. A low sterol content prevents ciliary membrane accumulation of the GPCR Smoothened (SMO), which functions as the transducer of the SHH signal. In this off-state, the SHH transcriptional effectors GLI2 and GLI3 are phosphorylated by cAMP dependent protein kinase (PKA) in the PC, which stimulates their partial proteolysis into transcriptional repressors ^12^. Pathway activation occurs upon SHH binding to PTCH to stimulate its internalization and degradation ^13^. Removal of PTCH from ciliary and plasma membranes allows for localized accumulation of sterols that bind SMO to stimulate its PC accumulation and signaling through two distinct effector routes. The first is a “noncanonical” SMO signal that activates Gαi heterotrimeric G proteins. This signal, which does not require SMO to be in the PC, controls transcription-independent SHH responses that influence lipid metabolism, Ca^2+^ release, and cytoskeletal regulation ^14–17^. The second effector route, referred to as the canonical SMO signal, requires SMO accumulation in the PC where it signals to block phosphorylation and partial proteolysis of GLI2/3. Full-length GLI2/3 proteins are then activated to induce the downstream transcriptional response ^12,14,18^. Current models suggest that accumulation and activation of SMO in the PC results from direct cholesterol binding to sites in its extracellular cysteine rich domain (CRD) and 7-transmembrane (7TM) bundle ^11,19–21^. Lipids including oxysterols, fatty acids, and phospholipids have also been reported to bind to sites in the CRD, 7TM bundle, and/or intracellular (IC) loops to promote SMO activation ^20,22–25^. Although the identity of the natural ligand and location of its primary orthosteric binding pocket remain topics of debate, most reports suggest that cholesterol binding to the SMO CRD and 7TM domains is crucial for signal induction ^11,19,20,22,24,26^. Accordingly, mutation of the residues that coordinate cholesterol binding disrupt SHH-mediated SMO activation ^11,20,24,27^.

Notably, many GPCRs have additional ligand binding pockets that are spatially distinct from the orthosteric site. These pockets are referred to as allosteric sites and, when occupied by allosteric ligands, allow for enhancement or suppression of orthosteric ligand-induced signaling activity ^28^. Consistent with this biology, we previously identified arachidonic acid (AA) as a feed-forward allosteric enhancer of SHH-activated SMO signaling ^15^. AA production occurs in response to noncanonical SMO signaling to activate Gαi, which stimulates cytosolic phospholipase A2α (cPLA2α) at the ciliary base to cleave arachidonate-containing phospholipids ^15^. Localized, SHH-stimulated synthesis of AA by cPLA2α at the PC serves two purposes. First, it leads to production of prostaglandin E2 (PGE2), which activates ciliary E-type prostanoid receptor 4 (EP4) to promote anterograde intraflagellar transport (IFT) for PC length stability and signaling competency ^29^. Second, it enhances SMO ciliary accumulation for optimal communication with GLI2/3 (Figure 1A). Inhibition of AA production reduces the amplitude of the SHH-induced signal response, suggesting that AA may function as a feed-forward allosteric enhancer of canonical SMO signaling ^15^.

**Figure 1.**
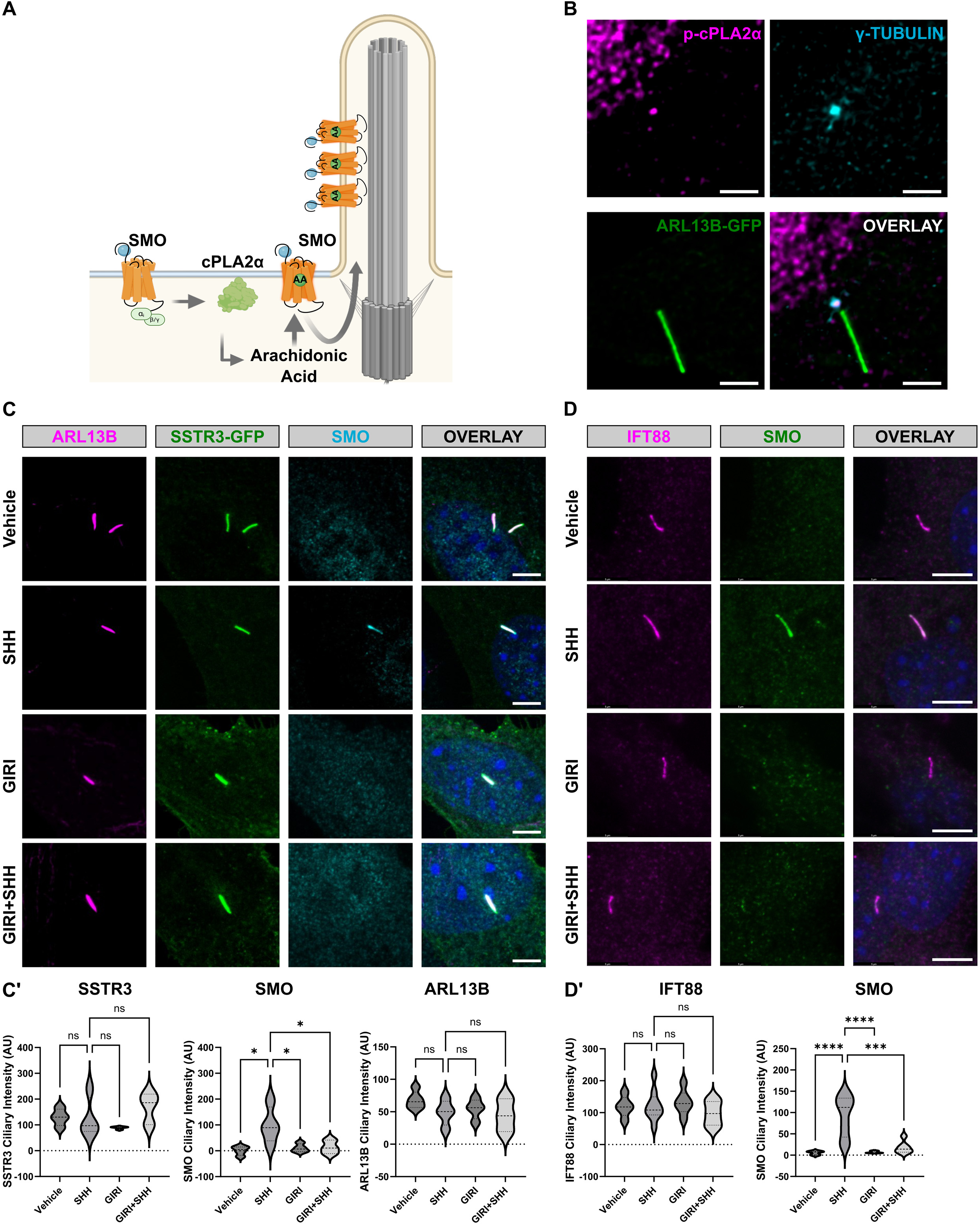
Specificity of cPLA2α contributions to ciliary trafficking. (A) SMO signal induction by a CRD-binding agonist (blue) activates Gαiβγ to stimulate cPLA2α production of arachidonic acid (AA). AA is proposed to interact with the 7TM domain of active SMO to enhance its enrichment in the PC for increased signaling to GLI2/3 transcriptional effectors (pink). (B) A fraction of endogenous phospho-cPLA2α (magenta) localizes to the base of the PC (marked by γ-tubulin, cyan) in NIH-3T3 cells. ARL13B-GFP (green) marks the ciliary axoneme. Scale bar = 2 μm. (C) NIH-3T3 cells expressing SSTR3-GFP were treated with vehicle (DMSO), GIRI (2μM), and control or SHH conditioned media as indicated for 18h. The ciliary axoneme is marked with ARL13B (magenta). Endogenous SMO is cyan. DAPI is blue. Scale bar = 5 μm. (C’) Quantification of ciliary signal intensity for SSTR3-GFP, SMO, and ARL13B in cells treated with control or SHH conditioned media plus 2 µM GIRI or vehicle control. The experiment was performed twice with 30 cilia imaged per condition per experiment and all data were pooled. (D) NIH-3T3 cells were treated with control or SHH conditioned media in the presence of 2 µM GIRI or vehicle. IFT88 is shown in magenta, SMO is green, DAPI (blue) marks nuclei. Scale bar = 5 μm. (D’) Quantification of ciliary signal intensity for IFT88 and SMO in cells treated with control or SHH conditioned media, 2 µM GIRI, or vehicle control. The experiment was performed twice with ≥30 cilia imaged per condition per experiment and all data were pooled. For all graphs, significance was determined using a one-way ANOVA. Significance is indicated as follows: *<0.05, **<0.01, ***<0.001, ****<0.0001, and ns, p > 0.05. Data are represented as mean ± SD.

Multiple reports have provided structural evidence and *in vitro* functional data supporting that SMO allostery enhances signal output ^11,15,22,24,25^. However, *in vivo* evidence for allosteric regulation of SMO signaling has remained elusive. Herein, we interrogate SMO allostery *in vitro* and *in vivo* to define the precise contribution of AA to SMO signaling activity during tissue morphogenesis. We show that AA enhances SMO signaling induced by TM-binding cholesterol and oxysterols, map a specific AA binding site at the top of the 7TM bundle, and identify an essential anchoring residue for AA in extracellular loop 2 (EC2). A single amino acid change at this site specifically displaces AA and blunts SMO signaling without compromising the structural integrity of the binding pocket. We introduce a novel knockin mouse model for reduced AA binding and show that homozygous mutation of the AA anchoring residue leads to high perinatal lethality. Neurodevelopment occurs normally in these animals, but cardiopulmonary development is disrupted. Altogether, these results suggest that AA allosterically enhances SMO signaling and that this functionality contributes to *in vivo* pathway activity during embryonic development in an organ-specific manner.

## Results

### AA specifically enhances SHH-stimulated SMO ciliary enrichment

The active phosphorylated form of cPLA2α localizes to the base of ARL13B-marked primary cilia of SHH-responsive NIH-3T3 cells, where it overlaps with the centrosome marker γ-tubulin (Figure 1B) ^15^. Enrichment of cPLA2α at the PC base raises the possibility that localized production of AA might impact ciliary enrichment of proteins other than SMO. To test this hypothesis, we expressed a green fluorescent protein (GFP)-tagged version of the ciliary GPCR Somatostatin Receptor 3 (SSTR3) in NIH-3T3 cells, and then monitored PC enrichment of SSTR3-GFP and endogenous SMO following treatment with control or SHH conditioned media and the cPLA2α inhibitor Giripladib (GIRI) ^30^. Consistent with our previous reports, GIRI treatment significantly reduced SMO ciliary enrichment in SHH-stimulated cells (Figure 1C-C’, cyan) ^15,29^. GIRI treatment did not impact SSTR3-GFP localization, indicating that AA production is unlikely to be a general modulator of GPCR ciliary translocation (Figure 1C-C’, green). PC enrichment of the GTPase ARL13B and the intraflagellar transport protein IFT88 was also unaffected by GIRI treatment, suggesting that cPLA2α is not a general modulator of PC protein trafficking and that the effect may be specific to SMO (Figure 1C-D’).

### AA enhances sterol-stimulated SMO signaling

SMO signaling can be induced by direct binding of cholesterol or oxysterols to distinct ligand binding pockets in its CRD and 7TM domains ^20–22,26^. We previously demonstrated that AA synergizes with the CRD-binding agonist 20(S)-hydroxycholesterol to augment SMO ciliary accumulation for enhanced signaling to GLI ^15^. AA does not induce a transcriptional response in SHH-responsive Light II reporter cells when provided in the absence of oxysterols, suggesting that AA might function as an allosteric activator of oxysterol or sterol-bound SMO (Figure 2A) ^15,31^. Thus, we tested whether AA would enhance pathway activation by other SMO-binding oxysterols, including CRD-binding 7β,27-dihydroxycholesterol (7β,27-DHC) and 24(S),25 epoxycholesterol (24,25-EpCHO), which influences SMO signaling through the CRD, 7TM bundle, and intracellular (IC) loops (Figure 2B) ^22,25^. SHH Light II cells were treated with 7β,27-DHC or 24,25-EpCHO alone or with increasing concentrations of AA, and normalized reporter activity was determined. Although both oxysterols induced a reporter response over baseline, induction was modest compared to the response induced by incubation with SHH-containing conditioned media (Figure 2C, column 1 vs. 2 and 6). Co-stimulation with increasing amounts of AA boosted reporter induction by CRD-binding 7β,27-DHC (Figure 2C, columns 3-5) and multisite-binding 24,25-EpCHO (columns 7-9). Notably, AA augmented 24,25-EpCHO-induced SHH Light II reporter stimulation similarly to SHH conditioned media (Figure 2C, columns 1 vs. 9). Enhancement of 7β27-DHC activity, which binds exclusively to the CRD, was comparatively modest (Figure 2B-C, columns 5 vs. 9).

**Figure 2.**
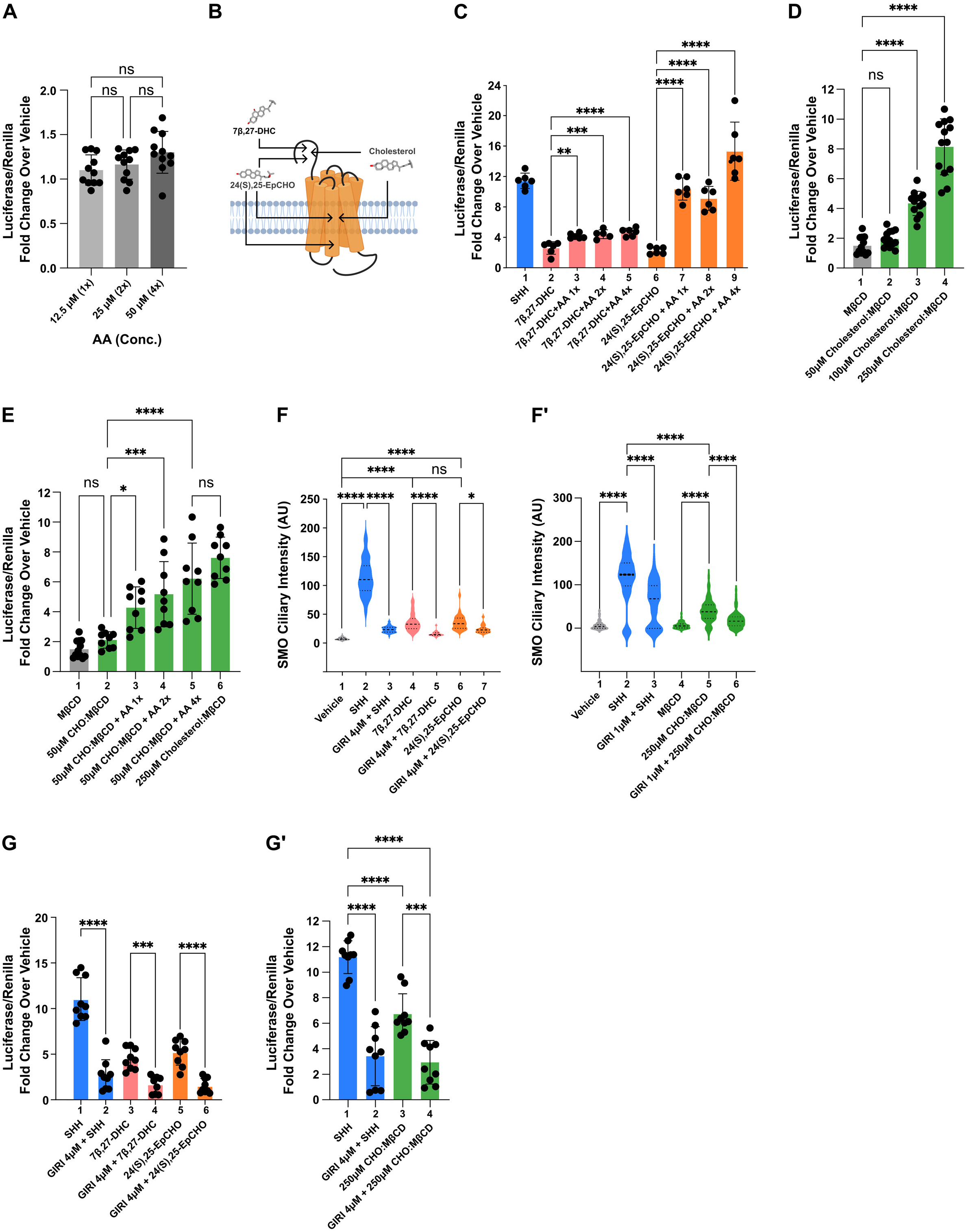
AA enhances sterol-mediated SMO activation. (A) SHH Light II cells were treated with the indicated concentrations of AA. The average Luciferase/Renilla value is shown as fold change over vehicle. The experiment was repeated three times with 3-4 technical replicates per experiment and all data were pooled. (B) A diagram of reported binding regions for oxysterols (7β,27-DHC and 24(S),25-EpCHO) and cholesterol (CHO) on SMO. (C) SHH Light II cells were treated with control or SHH conditioned media, and 15 µM 7β,27-DHC or 24(S),25-EpCHO alone or in combination with increasing concentrations of AA (1x=12.5 μM). The average Luciferase/Renilla value is shown as fold change over vehicle. The experiment was repeated three times with 2-3 technical replicates per experiment and all data were pooled. (D) Activation of SHH pathway in response to increasing CHO:MꞵCD (1:10) was measured in SHH Light II cells. The average fold change was calculated from three independent experiments, all done in triplicate. All data were pooled. (E) SHH Light II cells were treated with 50 μM CHO:MꞵCD plus vehicle or increasing concentrations of arachidonic acid (1x=12.5 μM). The maximum response is shown using 250µM CHO:MβCD. The average fold change was calculated from three independent experiments done in triplicate. All data were pooled. (F-F’) Quantification of SMO ciliary signal intensity in NIH-3T3 cells. (F) Cells were treated with DMSO, 30 µM 7β,27-DHC or 24(S),25-EpCHO following pretreatment with 4 µM GIRI or vehicle. (F’) Cells were treated with MꞵCD or CHO:MꞵCD (250 µM) plus control or SHH conditioned media following pretreatment with 1 µM GIRI or vehicle control. Lower concentrations of GIRI were used when combined with MβCD to prevent cells sloughing off the dish. Experiments were repeated twice with ≥75 cilia imaged per condition per experiment and all data pooled. (G-G’) cPLA2α is required downstream of SHH and SMO sterol agonists for maximal downstream transcriptional activation. SHH Light II cells were incubated with vehicle, 30 µM 7β,27-DHC, 30 µM 24(S),25-EpCHO, 250 µM CHO-MβCD (G’) and control or SHH conditioned media following vehicle or 4 µM GIRI pretreatment. Luciferase expression was normalized to Renilla. The average fold change over vehicle control was calculated from three independent experiments done in triplicate and all data were pooled. Error bars indicate SD. Statistical significance was determined using one-way ANOVA and indicated as follows: *<0.05, **<0.01, ***<0.001, ****<0.0001, and ns, p > 0.05.

Structural studies have revealed that, like oxysterols, cholesterol can associate through ligand binding pockets in the CRD and deep within the 7TM bundle of SMO to promote its ciliary enrichment for signaling to GLI2/3 (Figure 2B) ^22,32,33^. Consistent with these reports, treatment of SHH Light II cells with increasing concentrations of cholesterol-loaded methyl-β-cyclodextrin (50-250 μM CHO:MβCD) ^34^ increased GLI reporter activity in a dose-dependent manner (Figure 2D). To determine whether AA would impact CHO:MβCD activation of SMO, we titrated increasing concentrations of AA into SHH Light II cell culture media along with 50 µM CHO:MβCD. AA increased GLI reporter induction in a dose-dependent manner, raising the 50 µM CHO:MβCD response to a level comparable to that of high-dose CHO:MβCD (250 µM, Figure 2E, columns 2-5 vs. 6).

Having established that addition of exogenous AA enhanced SMO signal induction by CRD- and 7TM-binding sterols and oxysterols, we next sought to determine whether 7β,27-DHC, 24,25-EpCHO, or CHO:MβCD required activity of the AA-producing enzyme cPLA2α for maximal signaling. To evaluate upstream pathway activation, we quantified SMO ciliary enrichment following oxysterol or cholesterol stimulation in cells treated with vehicle or GIRI. The ability of SHH or activating sterols to induce robust SMO ciliary accumulation was significantly reduced by GIRI treatment (Figure 2F-F’ and Supplementary Figure 1). Downstream signaling to GLI was similarly attenuated as demonstrated by significantly reduced SHH-, 7β,27-DHC-, 24,25-EpCHO-, and CHO-MβCD-stimulated GLI reporter activity in GIRI treated SHH Light II cells (Figure 2G-G’). Taken together with the results presented above, these experiments demonstrate that cPLA2α-produced AA enhances both sterol and oxysterol-stimulated SMO ciliary accumulation and downstream signaling.

### AA is predicted to bind at the top of the SMO 7TM bundle

Having established that AA enhances signaling by sterol-stimulated SMO, we next sought to determine whether signal augmentation resulted from AA binding to a specific pocket in the CRD, EC loops, or 7TM domains. To identify a candidate binding site for AA, molecular docking experiments were performed using Maestro software (Schrödinger Release 2019-3) with receptor grids based upon published crystal structures of sterol- and cyclopamine-bound *Xenopus* and human SMO proteins ^24,35,36^. Grids were validated by redocking cholesterol onto the CRD and the inverse agonist cyclopamine onto the upper TM/EC loop pocket (Figure 3A). The high degree of overlap between native and docked binding molecules supported feasibility of using *in silico* docking to predict how AA might bind SMO (Figure 3A green vs. yellow). Experimental *in silico* docking predicted AA binding near the top of the SMO 7TM domain in a pocket that significantly overlapped with the cyclopamine and SAG binding site (Figure 3B-D) ^32^. The high degree of predicted overlap between AA and cyclopamine is consistent with our previously published observation that AA can displace Bodipy-cyclopamine from SMO in cell-based binding assays ^15^. Although cyclopamine and AA were predicted to occupy similar space in the upper 7TM/EC pocket, they were predicted to anchor through distinct residues in the EC loops and 7TM helices (Figure 3B-C’). Cyclopamine anchors through a hydroxyl group that forms a hydrogen bond with the side chain of E491 in TM helix 7 (Figure 3C). In its most favorable predicted pose, the hydroxyl group of AA is deprotonated to form an O^−^ anion that facilitates a salt bridge with the α-amino group of K368 in EC loop 2 (Figure 3C’). Intriguingly, the predicted topology of AA association with K368 of *Xenopus* SMO positions the fatty acid in a binding pocket that is predicted to be offset from the deep 7TM cholesterol pocket, suggesting that the two molecules may be able to bind simultaneously (Figure 3D). Notably, *Xenopus* SMO K368 corresponds to K395 of human SMO, which is reported to contribute to binding of several small molecule SMO inhibitors ^22,37^. Alignment of SMO protein sequence revealed high conservation of the predicted AA anchoring lysine across most vertebrate SMO proteins, further suggesting physiological significance of the residue (Figure 3E).

**Figure 3.**
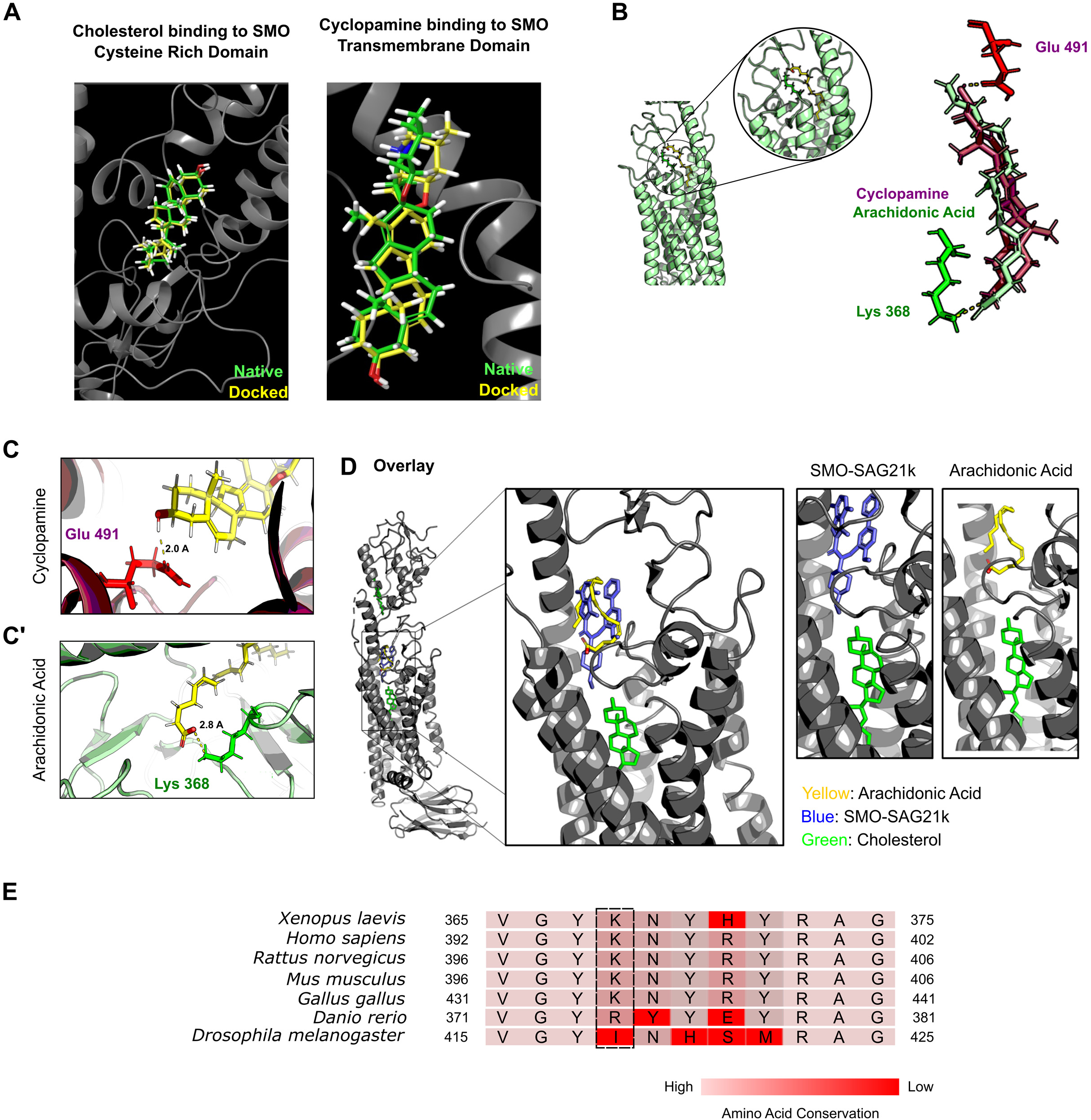
*In silico* docking predicts an AA binding site on SMO. (A) *In silico* docking (yellow) of cholesterol recapitulates the native (green) binding pose observed in the crystal structure of cholesterol bound to the human SMO extracellular CRD (PDB ID: 5L7D). Docking of cyclopamine occupies the same pocket with a similar binding pose as in the crystal structure of cyclopamine bound to the *Xenopus* SMO transmembrane domain (PDB ID: 6D32). (B) *In silico* docking of cyclopamine and AA onto *Xenopus* SMO using Maestro software predicts cyclopamine and AA bind through an overlapping pocket near the top of the 7TM of SMO. (C) Cyclopamine is anchored by *Xenopus* SMO E491 (E522 in murine SMO) in transmembrane helix 7. (C’) AA is predicted to form a salt bridge with *Xenopus* SMO K368 (murine SMO K399) in EC2. (D) *In silico* docking predicts that the AA (yellow) binding pocket overlaps with the SAG (blue) pocket, but not with the deep 7TM cholesterol (green) binding pocket of the murine SMO structure (PDB ID: 6O3C). (E) Alignment of EC2 sequence from SMO proteins. The predicted AA anchoring residue is conserved across most vertebrate species (dotted box).

To test whether AA association with SMO required the conserved lysine, an alanine substitution was introduced at the corresponding residue in murine SMO-YFP (K399A), and then the ability of AA to confer thermal stability through direct binding to wild-type (WT) and K399A SMO-YFP proteins was evaluated ^38^. Cyclopamine binding deficient E522A SMO-YFP was used as control ^33^. Membrane fractions from HEK293T cells expressing WT, E522A, or K399A SMO-YFP proteins were incubated with AA, cyclopamine, or vehicle control for 1 hour at 34°C prior to shifting to a higher temperature. Following thermal shift, membrane fractions were evaluated by SDS-PAGE and western blot, and densitometry analyses were performed to quantify SMO-YFP signal intensities at the increased temperatures. Consistent with direct binding of cyclopamine and AA to SMO-YFP, thermal stability of the WT SMO-YFP protein was enhanced by incubation with either molecule. Cyclopamine increased the Tm by ∼9°C and AA increased the Tm by ∼5.7°C (Figure 4A and B). Whereas K399A mutation did not significantly alter the Tm shift induced by cyclopamine, it blocked the ability of AA to stabilize SMO, supporting that K399 contributes to AA-SMO binding (Figure 4A vs. A’ and B vs. B’). Maintenance of the cyclopamine-induced thermal shift of SMO^K399A^-YFP suggests that failure of AA to stabilize the protein resulted from a specific attenuation of AA binding, and not a K399A-induced protein folding abnormality (Figure 4B’). Reciprocal results were obtained with the cyclopamine binding deficient E522A mutant. AA enhanced the thermal stability of SMO^E522A^-YFP by approximately 4.5°C, which is comparable to the shift observed for the WT protein incubated with AA (Figure 4A vs. A”). Conversely, thermal stability resulting from addition of cyclopamine was blunted from an increase of ∼9.0°C for the WT protein to an increase of ∼4.8°C for SMO^E522A^-YFP (Figure 4B vs. B”). Taken together, these results suggest that K399 is important for AA association with SMO.

**Figure 4.**
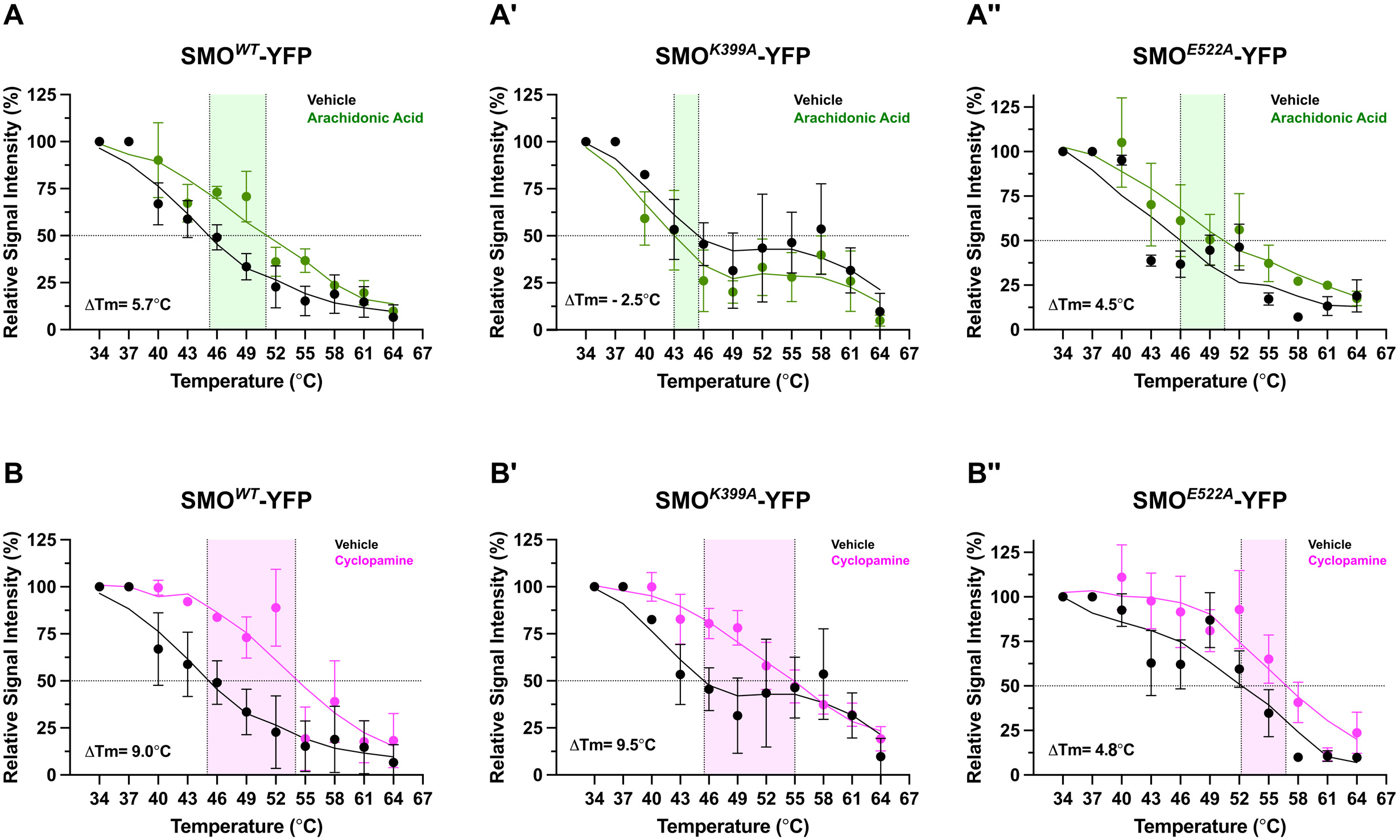
K399 facilitates AA binding to murine SMO. (A-B”) Ligand-induced thermal stability changes were evaluated for wild-type (WT), K399A (AA binding mutant) and E522A (cyclopamine binding mutant) SMO-YFP proteins. Membrane fractions from cells expressing the indicated SMO-YFP proteins were incubated with vehicle, 12.5 μM AA, or 10 µM cyclopamine and then incubated at the indicated temperatures. Thermal stability was assessed by western blot. Signal intensity relative to the 34°C start point (set to 100%) is shown. The experiment was repeated twice for SMO^E522A^-YFP + AA and three times for all other conditions. All data were pooled and shown on the summary graphs. Error bars indicate SEM.

### AA binding allosterically activates SMO signaling

Having determined that AA is unlikely to bind SMO^K399A^ as efficiently as it does the WT protein, we next sought to determine whether SMO^K399A^ demonstrated attenuated ligand-induced PC enrichment or signaling activity. Because transient high-level over-expression of SMO leads to ligand-independent ciliary localization, we knocked out endogenous *Smo* in Flp-In competent NIH-3T3 cells using CRISPR/Cas9 and then introduced WT SMO, K399A or E522A SMO-YFP cDNAs. We sorted YFP-positive cells and selected clones expressing WT, SMO^K399A^ and SMO^E522A^ -YFP proteins at comparable low levels (Figure 5A). We confirmed that SMO^K399A^ and SMO^E522A^ mutant proteins reached the cell surface like WT SMO-YFP, and then used these validated cell lines to test SMO ciliary entry and downstream signaling activity (Figure 5B and Supplementary Figure 2A-B). WT SMO-YFP accumulated in cilia following treatment with activating ligands SHH and SAG and the inverse agonist cyclopamine, which promotes SMO ciliary enrichment despite blocking its downstream signaling to GLI2/3 (Figure 5C-C’ and Supplementary Figure 2C) ^39^. We previously demonstrated that SHH, but not SAG, requires cPLA2α-mediated production of AA to stimulate robust SMO ciliary enrichment ^15^. Accordingly, AA binding-deficient SMO^K399A^-YFP showed significantly reduced PC enrichment in response to SHH stimulation but maintained its ability to enter cilia following treatment with SAG or cyclopamine, which bind to the same pocket (Figures 3D and 5C-C’ and Supplementary Figure 2C) ^33^. Cyclopamine treatment did not effectively induce ciliary enrichment of the E522A mutant protein, but SHH and SAG exposure did (Figure 5C-C’ and Supplementary Figure 2C). Importantly, diminished SHH-stimulated ciliary enrichment of SMO^K399A^-YFP correlated with reduced downstream signaling activity. Whereas SHH exposure stimulated GLI1 protein production in cells expressing WT or E522A SMO-YFP proteins, SHH-stimulated GLI1 was nearly undetectable in SMO^K399A^-YFP Flp-In cells (Figure 5D, lanes 3-4). Thus, direct AA-SMO binding likely contributes to SMO ciliary enrichment and signaling in SHH-stimulated cells.

**Figure 5.**
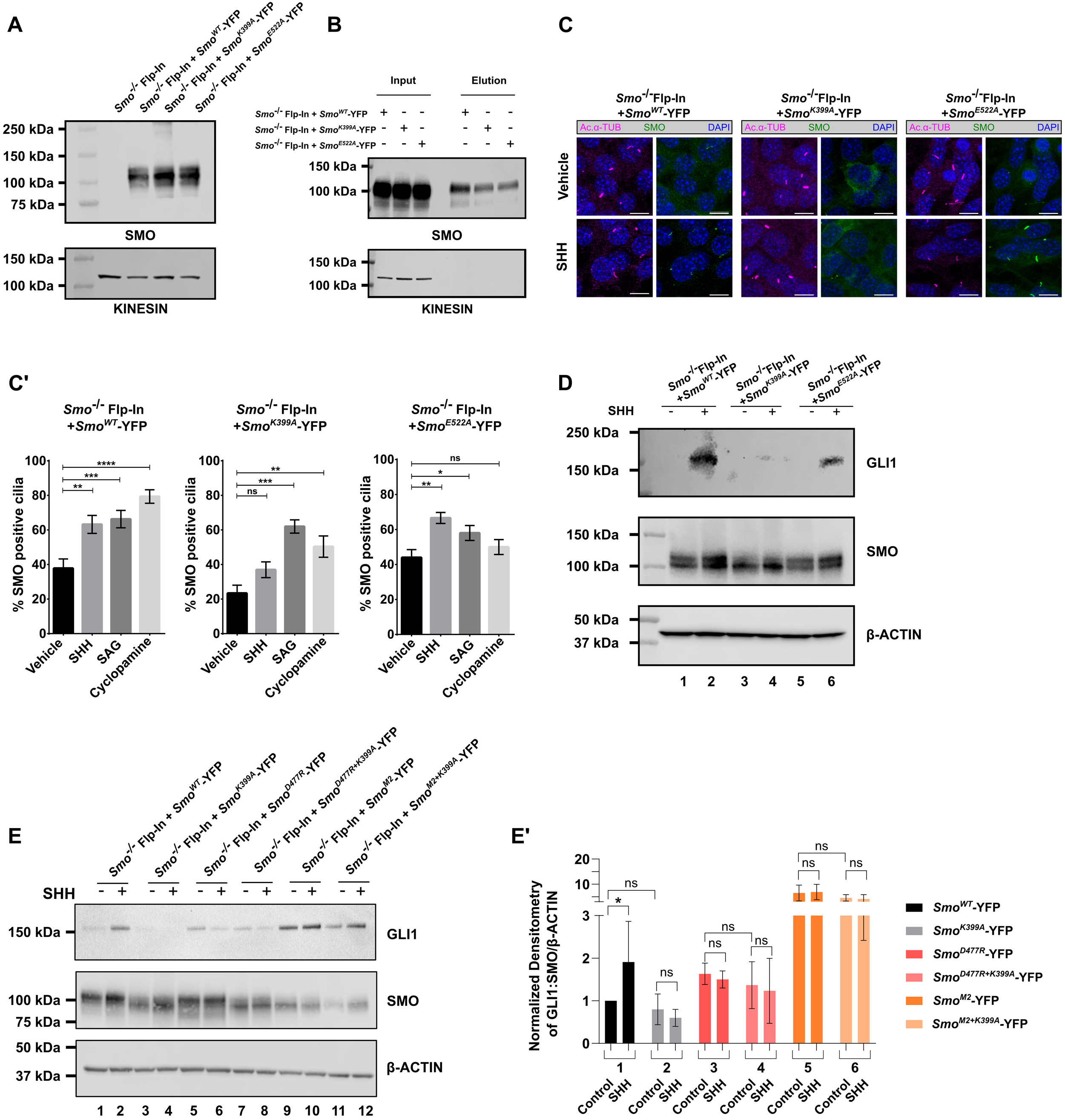
SMO^K399A^ shows reduced ligand responsiveness. (A) *Smo^−/−^* Flp-In-NIH-3T3 cells were stably transfected with *Smo^WT^-YFP*, *Smo^K399A^-YFP*, or *Smo^E522A^-YFP* cDNAs. Protein levels in cell lysates were analyzed by western blot. Kinesin is the loading control. (B) Cell surface biotinylation was performed on *Smo^−/−^* cells stably expressing YFP-tagged *Smo^WT^*, *Smo^K399A^* or *Smo^E522A^*. Biotinylated proteins were collected on streptavidin beads and analyzed by western blot. Kinesin marks the cytoplasmic input. (C) SHH-stimulated SMO ciliary translocation in the indicated cell lines was evaluated by confocal microscopy. The ciliary axoneme is marked by acetylated α-tubulin (magenta) and SMO is shown in green. DAPI (blue) marks nuclei. Scale bar = 5 μm. (C’) Flp-In-NIH-3T3 cells expressing the indicated SMO constructs were treated with control or SHH-containing conditioned media and vehicle, SAG, or cyclopamine, and then SMO ciliary localization was quantified. Results are presented as percent cilia with positive SMO signal. The experiment was performed twice with ≥50 cells counted for each condition. All data were pooled. (D-E) GLI1 and SMO protein levels were analyzed in lysates from control or SHH conditioned media treated *Smo^−/−^* Flp-In-NIH-3T3 cells stably transfected with the indicated *Smo-YFP* cDNAs. ꞵ-actin is the loading control. Experiments were performed three times. Representative blots are shown. (E’) Densitometry analysis of GLI1, SMO, and β-actin was performed. GLI1:SMO ratios relative to β-actin are shown. The graph shows pooled data from 3 independent experiments. For all panels, significance was determined using a one-way ANOVA. Significance is indicated as follows: *<0.05, **<0.01, ***<0.001, ****<0.0001, and ns, p > 0.05.

Based on its predicted binding site at the top of the 7TM domain, we hypothesized that AA may function by “capping” the SMO 7TM core to stabilize sterol binding deep in the 7TM pocket. If this is the mechanism by which AA enhances sterol-mediated SMO activation, we reasoned that the functional consequence of the K399A mutation might be mitigated by introducing the activating D477R mutation, which is proposed to stabilize 7TM sterol binding by a similar mechanism ^22,37^. The oncogenic M2 modification, which stabilizes a ligand-independent active TM conformation, was also evaluated ^40^. Compound and single amino acid SMO-YFP mutants were stably expressed in Flp-In competent *Smo^−/−^* NIH-3T3 cells as described above, and SMO N-linked glycan modifications were analyzed to test for ER retention of single and compound mutant proteins. PNGase and EndoH treatment revealed that SMO^K399A^ and SMO^D477R^ single and compound mutant variants produced EndoH-resistant fractions, consistent with normal trafficking out of the ER (Supplementary Figure 2A) ^41^. Accordingly, both variants reached the plasma membrane, as indicated by cell surface biotinylation (Supplementary Figure 2B). SMO^M2^ uses an unconventional secretory route that bypasses the Golgi where complex glycans are added ^42^. Accordingly, we did not detect EndoH-resistant N-linked glycosylation for either SMO^M2^ or SMO^M2+K399A^ (Supplementary Figure 2A). Reduced cell surface biotinylation of SMO^M2^ variants compared to controls was also noted (Supplementary Figure 2B).

To evaluate signaling competency of each of the single and compound mutant SMO variants, cells were exposed to control or SHH conditioned media, and induction of endogenous GLI1 was evaluated by western blot. Given the differing proteins levels of the compound mutant SMO variants, GLI1:SMO ratios were determined by densitometry, and then normalized to the β-actin loading control. Expression of either SMO^D477R^-YFP or SMO^M2^-YFP led to increased baseline induction of GLI1 that was not further elevated upon SHH exposure (Figure 5E, lanes 1-2 vs. 5-6 and 9-10 and E’). Intriguingly, constitutive SMO signaling activity that was conferred by either D477R or M2 mutations was unaltered by the introduction of the K399A mutation as GLI:SMO ratios did not change (Figure 5E’ columns 3 vs. 4 and 5 vs. 6). These results suggest that activating mutations that promote SMO signaling through 7TM capping or conformational shifts do not require AA binding to the 7TM domain.

### Allostery promotes in vivo SMO signaling

To interrogate whether allostery at the 7TM binding pocket is required for *in vivo* SMO signaling, we used CRISPR/Cas9 to generate K399A knockin mice (Supplementary Figure 3A) and evaluated control and *Smo^K399A/K399A^* mice for phenotypic indicators of altered developmental patterning. Whereas *Smo^K399A/K399A^* mice were present at expected Mendelian ratios as embryos, they were underrepresented following birth (Figure 6A). Reduced numbers of *Smo^K399A/K399A^* mice were evident by postnatal day P5 and worsened by weaning at P21 (Figure 6A-B). Whereas no survival or overt phenotypic differences were observed between *Smo^+/+^* and *Smo^K399A/+^* animals, *Smo^K399A/K399A^* mice that survived past birth were reduced in size and failed to thrive (Figure 6B-C’). Accordingly, computed tomography (CT) scans revealed the skulls of *Smo^K399A/K399A^* mice to be significantly smaller across several reference axes than *Smo^K399A/+^* littermate controls (Supplementary Figure 3B-B’). These results suggest that SMO allostery contributes to SHH signaling during development.

**Figure 6.**
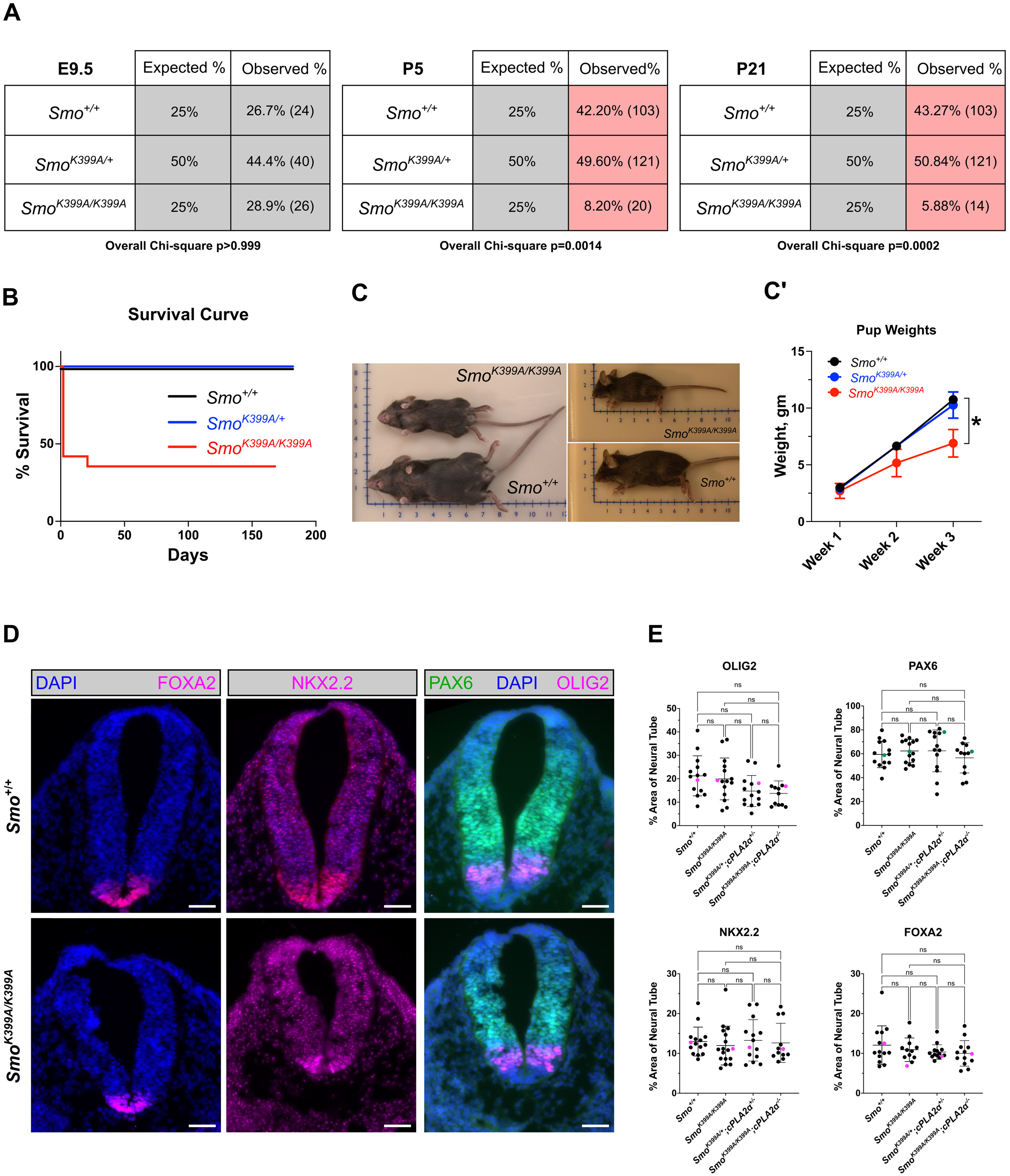
*Smo^K399A/K399A^*knockin mice fail to thrive. (A) *Smo^K399A/K399A^* knockin mice show normal Mendelian ratios at E9.5 and exhibit skewed Mendelian ratios at P5 and P21. Overall Chi-square values are shown for each timepoint. (B) A Kaplan-Meier survival curve shows reduced viability of *Smo^K399A/K399A^* mice compared to *Smo^K399A/+^* and *Smo^+/+^* controls. Litters were monitored weekly from P0. *n* ≥ 200 total mice analyzed. (C-C’) *Smo^K399A/K399A^* mice are reduced in size (C) and weight (C’) compared to controls. (D) Cardiac level sections of developing neural tubes of E9.5/25-29 somite stage embryos were stained for the floor plate marker FOXA2 and ventral progenitor markers NKX2.2, OLIG2, and PAX6. DAPI marks nuclei. Scale bar = 50μm. (E) Mean expression domain areas of the indicated progenitor markers were measured and normalized to overall neural tube area. At least 3 embryos per genotype with 3-5 sections per embryo were analyzed. Data are represented as mean ± SD. For C’ and E, statistical significance was calculated using a Student’s t-test between *Smo^+/+^* and *Smo^K399A/K399A^* and is indicated by *p < 0.01 and ns = not significant.

We reasoned that if poor viability of knockin mice resulted from attenuation of SMO signaling stemming from compromised AA allostery, combining SMO^K399A^ mutation with decreased AA production would likely enhance the observed phenotype. Thus, we obtained *cPLA2α^−/−^* mice ^43^ to remove activity of the AA-producing phospholipase that is induced downstream of SHH ^15^. Due to compensation by other phospholipases, *cPLA2α^−/−^* mice are viable, but females exhibit reduced fertility ^43^. Strikingly, removal of one or both copies of *cPla2α* in the *Smo^K399A/K399A^* background significantly enhanced the perinatal lethality of the *Smo^K399A/K399A^* mice. Only two *Smo^K399A/K399A^;cPla2α^−/−^* and four *Smo^K399A/K399A^;cPla2α^+/−^* mice were recovered out of 268 total animals screened (Supplementary Table 1). All recovered compound mutants were male. Curiously, we did not observe worsening of the *Smo^K399A/K399A^* size reduction phenotype in the surviving *Smo^K399A/K399A^;cPla2α^−/−^* mice, suggesting that they may represent phenotypic “escapers” with compensatory signaling activity (Supplementary Figure 3C). Taken together with *in vitro* data that SHH-mediated induction of cPLA2α stimulates production of AA that anchors through the K399 residue, these results suggest that AA enhances *in vivo* SMO signaling.

Because SHH is a key regulator of multiple organs and tissues during embryonic development ^44^, identifying the cause of reduced viability of *Smo^K399A/K399A^* mutants necessitated empirical determination. We began by evaluating the nervous system because, during neural tube patterning, SHH activates distinct transcriptional programs to define specific neural progenitor cell populations in the ventral neural tube ^45^. We analyzed neural tubes of *Smo^+/+^*, *Smo^K399A/+^;cPla2α^+/−^*, *Smo^K399A/K399A^*, and *Smo^K399A/K399A^;cPla2α^−/−^* E9.5/25-29 somite stage embryos for induction of the floor plate marker *FoxA2*, ventral fate markers *Nkx2.2* and *Olig2* and intermediate fate marker *Pax6* (Figure 6D-E and Supplementary Figure 3D). Despite the observed viability defect, ventral progenitor domain establishment in *Smo^K399A/K399A^* embryos was comparable to *Smo^+/+^* controls (Figure 6D-E). Ventral neural tube progenitor domain specification was similarly unaltered in *Smo^K399A/+^;cPla2α^+/−^* and *Smo^K399A/K399A^;cPla2α^−/−^* E9.5/25-29 somite stage embryos (Figure 6E and Supplementary Figure 3D). These results suggest that the early stages of nervous system development proceed normally in mice harboring the *Smo^K399A^* allele.

To determine whether neuronal patterning defects arose at later stages of development, we analyzed cerebella of *Smo^K399A/+^* and *Smo^K399A/K399A^* adult mice. No significant differences were detected in the ratios of cerebellar to total brain volumes between control and *Smo^K399A/K399A^* mice (Supplementary Figure 3E-E’). Taken together with the results presented above, cerebellar analyses suggest that SMO allostery through the K399 anchor is not essential for neuronal cell fate specification and that reduced viability of *Smo^K399A/K399A^* mice was not due to compromised neurodevelopment.

In addition to its contributions to nervous system development, SHH also plays important roles during patterning of the heart and lungs ^3,46–48^. During heart development, SHH drives heterochronic gene expression to ensure proper timing of second heart field (SHF) progenitor differentiation and morphogenesis ^4^. In the lung, SHH signaling is active during three key stages of lung morphogenesis including initial budding of the lung from the foregut around E9.5, branching morphogenesis near E11.5, and alveolar remodeling at birth ^3,6,49^. We reasoned that the high perinatal lethality associated with SMO^K399A^ mutation could result from compromised cardiopulmonary development, as suggested by observed “gasping” and a cyanotic appearance of P0 *Smo^K399A/K399A^* mice (Figure 7A). To probe for indicators of altered heart or lung development in *Smo^K399A/K399A^* embryos, we performed bulk RNAseq analysis on E9.5 *Smo^K399A/K399A^* and *Smo^+/+^* littermate control embryos (Supplementary Figure 4A-F’). Among the top pathways upregulated in *Smo^K399A/K399A^* embryos were heart development, TGFβ signaling, and prostaglandin signaling, which is driven by AA metabolism. Top downregulated pathways included multiple genes related to PC function and ciliopathies (Supplementary Figure 4A-F’).

**Figure 7.**
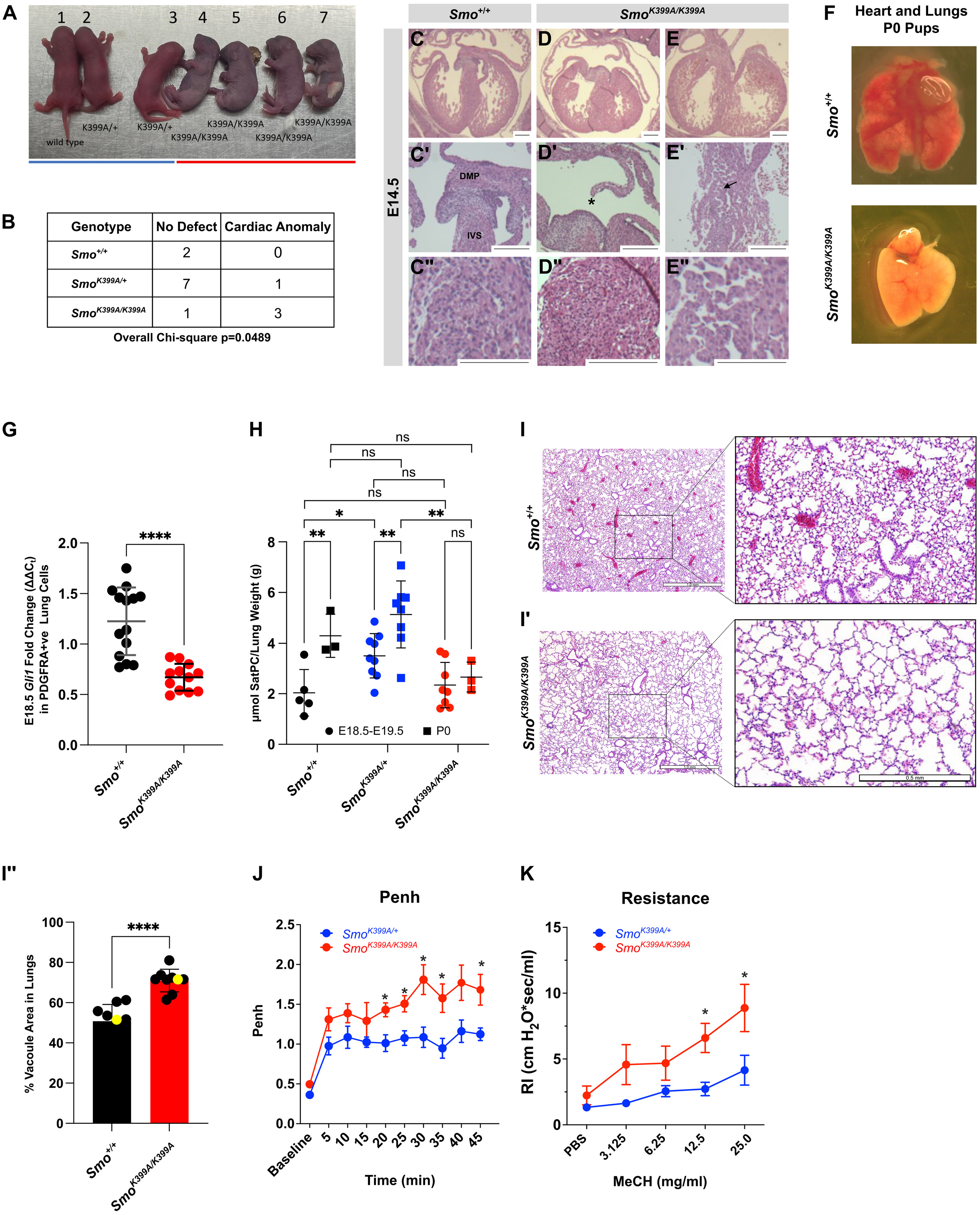
*Smo^K399A/K399A^*mice have cardiopulmonary defects. (A) A P0 litter shows the cyanotic appearance of *Smo^K399A/K399A^* mice compared to *Smo^+/+^* and *Smo^K399A/+^* littermates. (B) Overall Chi-square analysis correlating the effect of *Smo^K399A/K399A^* mutation with cardiac mispatterning. (C-E”) Transverse cardiac sections of E14.5 embryos were stained with Hematoxylin/Eosin (H&E). Whole heart sections (C, D, E) and higher magnifications (C’-E”) of *Smo^+/+^* and *Smo^K399A/K399A^* embryos reveal AVSD (C’-D’) and IVS (C”-E”) patterning defects (arrow). DMP = Dorsal Mesenchymal Protrusion, IVS = Interventricular Septum, * = Atrioventricular Septal Defect. Scale bar = 200µm. (F) Hearts and lungs of P0 *Smo^+/+^* and *Smo^K399A/K399A^* mice are shown. (G) qRT-PCR analysis of *Gli1* expression in PDGFRA^+^ cells isolated from E18.5 lungs of *Smo^+/+^* and *Smo^K399A/K399A^* mutant mice is shown. Fold-change in expression was determined using the 2^−ΔΔCt^ method. Average fold change was calculated across 3 technical replicates each from 5 *Smo^+/+^* and 4 *Smo^K399A/K399A^* embryos. (H) Saturated phosphatidylcholine (SatPC) was measured in dissected embryonic (E18.5-E19.5) and P0 lungs. Total SatPC for each mouse was normalized to lung weight. At least 3 mice per genotype per timepoint were analyzed and all data were pooled. Error bars indicate SD. (I-I’) H&E stains of adult (P27) lung sections from *Smo^+/+^* and *Smo^K399A^*^/K399A^ mice are shown. Scale bar = 0.5 mm. (I”) Quantification of vacuole area in lung sections from *Smo^+/+^* and *Smo^K399A^*^/K399A^ animals. At least 2 randomly chosen 8x zoom regions of lung sections from 5 mice each were evaluated. The yellow dots indicate the sections shown in (I). Significance was calculated using a Student’s t-test and denoted as: *<0.05, **<0.01, ****<0.0001. Error bars indicate SD. (J) *Smo^K399A/+^* and *Smo^K399A/K399A^* mice were subjected to whole-body plethysmography. Enhanced pause (Penh) values were recorded over 45 minutes at 5-minute intervals. Results are plotted as maximal fold increase in Penh relative to baseline and expressed as mean ± SE, where *n*=6 mice per group. (K) Airway resistance was evaluated before and after exposure to the indicated concentrations of aerosolized methacholine (MeCH). * Difference from the *Smo^K399A/+^* mice, p< 0.05.

Given the established link between SHH signaling, ciliopathies, and congenital heart disease ^50–52^, we hypothesized that disruption of ciliary SMO signaling in *Smo^K399A/K399A^* embryos might lead to compromised heart development. To interrogate this, hearts of control and *Smo^K399A/K399A^* E14.5 embryos were subjected to histological examination. Atrioventricular septum defects (AVSD) were observed in 3 of 4 mutant embryos analyzed (Figure 7B and C-C’ compared to D-D’, asterisk). We also observed mispatterning of the interventricular septum (IVS) in *Smo^K399A/K399A^* embryos (Figure 7C’-C” compared to E’-E”, arrow). These defects are consistent with the known manifestations of reduced SHH signaling during cardiac progenitor field establishment ^4^. The observed increase in heart development and differentiation drivers including *Tbx5, Myocd,* and *Cyp26c1* observed in RNAseq data sets from E9.5 *Smo^K399A/K399A^* embryos are consistent with failed heterochronic differentiation control (Supplementary Figure 4A-B, D-E) ^4,53^.

Significant signal crosstalk occurs between the developing heart and lungs, suggesting that lung development might also be compromised in *Smo^K399A/K399A^* mice ^54^. Consistent with this hypothesis, lungs of P0 *Smo^K399A/K399A^* mice showed altered morphology compared to controls (Figure 7F). To evaluate SHH signaling activity in developing lungs, we purified SHH-responsive PDGFRα^+^ fibroblasts from lungs of E18.5 control and *Smo^K399A/K399A^* embryos and then quantified *Gli1* expression in the PDGFRα^+^ population by qPCR analysis ^55^. *Gli1* expression was significantly reduced in *Smo^K399A/K399A^* mutants compared to control, indicating compromised SHH activity in prenatal lungs (Figure 7G and Supplementary Figure 4G) ^56^. To determine whether reduced SHH signaling activity in PDGFRα+ fibroblasts at E18.5 led to alveolar maturation defects in *Smo^K399A/K399A^* neonates, saturated phosphatidylcholine (SatPC) was measured as an indicator of lung surfactant production in embryonic and P0 mice. Surfactant secretion from alveolar type II (AT2) cells at birth facilitates the transition to air breathing by lowering air-liquid surface tension and preventing alveolar collapse ^57^. Failure of AT2 cells to produce or secrete surfactant compromises this essential transition, which can result in gasping or inability to breathe at birth ^58^. Although the precise signals that stimulate surfactant secretion are not known, *in vitro* studies suggest that cPLA2α and AA signaling may be involved ^59,60^.

Accordingly, we found that while *Smo^+/+^* and *Smo^K399A/+^* mice increased surfactant SatPC levels at birth, *Smo^K399A/K399A^* mice, which we hypothesize have reduced SMO responsiveness to AA, did not show a SatPC shift (Figure 7H). Although *Smo^K399A/+^* animals showed elevated embryonic SatPC levels compared to *Smo^+/+^* control, SatPC showed a comparable fold increase over the E18.5-19.5 baseline at birth, suggesting that control of surfactant production and release is intact. We do not know the reason for the increased baseline SatPC levels in heterozygous animals but speculate that altered AA metabolism in *Smo^K399A/+^* animals may contribute.

Although most *Smo^K399A/K399A^* mice died during the perinatal period, those that survived to P21 typically reached adulthood (Figure 6B). To determine whether the surviving *Smo^K399A/K399A^* mice showed evidence of pulmonary mispatterning, we evaluated alveolar morphology in lung sections from control and surviving *Smo^K399A/K399A^* mice. Hematoxylin and Eosin (H&E) staining of adult lungs revealed alveolar simplification in *Smo^K399A/K399A^* lung sections, suggestive of reduced efficiency of gas exchange (Figure 7I-I”) ^6^. To evaluate the consequence of altered alveolar patterning in mutant mice, we performed unrestrained whole-body plethysmography (WBP) on control and surviving *Smo^K399A/K399A^* adults ^61^. WBP revealed that *Smo^K399A/K399A^* mice had significantly higher Penh scores compared to *Smo^K399A/+^* control animals, indicating increased airway hyperresponsiveness and altered lung function (Figure 7J). To test whether this physiological abnormality correlated with altered lung mechanics, *Smo^K399A/K399A^* and control animals were intubated and exposed to methacholine (MeCH), and airway resistance was measured ^62^. We noted significantly increased airway resistance in *Smo^K399A/K399A^* mice compared to *Smo^K399A/+^*, control animals, indicating higher susceptibility to bronchoconstriction (Figure 7K). Altogether with the observed cardiac phenotypes, neonatal and adult lung analyses suggest that *in vivo* SMO allostery contributes to optimal SHH signaling during cardiopulmonary development.

## Discussion

Allosteric regulation of receptor and enzymatic effector activity provides cells with the ability to tune the amplitude of a ligand-induced response to meet context-specific signaling requirements. This may be one mechanism by which morphogens, which signal in time- and concentration-dependent manners, pattern tissues with temporal and spatial precision. Consistent with this hypothesis, the signal transducer activated downstream of the SHH morphogen, the GPCR SMO, is proposed to be controlled through distinct orthosteric and allosteric ligand binding pockets ^11^. However, *in vivo* evidence for SMO allostery was lacking. The results presented here begin to close this knowledge gap by identifying AA as an allosteric enhancer of SMO signaling during cardiopulmonary development. We identified murine SMO K399 as an AA-anchoring residue and demonstrated that its mutation attenuates SHH-stimulated SMO ciliary enrichment, blunts downstream signaling, and triggers cardiopulmonary phenotypes in *Smo^K399A/K399A^* knockin mice.

Although SHH signaling is a known contributor to heart and lung development, its dysfunction is most linked to compromised neurodevelopment ^2,63^. As such, it was unanticipated that neural tube patterning and cerebellar development would be unaltered in *Smo^K399A/K399A^* mice and that these animals would instead present with cardiopulmonary mispatterning. One interpretation of this result is that SMO allostery is context-specific such that the developing heart and lungs need allosteric amplification of SMO signaling while the neural tube does not. Cell fate determination during neural tube patterning occurs in response to the combined action of opposing morphogen signals controlling robust gene regulatory networks that reinforce neuronal fate choices over time ^64^. Thus, the neural tube may have sufficient temporal and spatial feedback mechanisms in place to compensate for hypomorphic activity of the SMO^K399A^ mutant. By comparison, heart and lung development depends on strong bursts of SHH signaling activity at distinct developmental time points when crucial patterning decisions are made. In the heart, first and second heart fields are directed to cardiomyocyte fates as early as E7.25, but SHF commitment must be delayed until ∼E10.0 so the structures that are necessary for pulmonary circulation specify at the correct time and place ^65,66^. SHH instructs this precise timing such that delay or reduction of signal activation leads to cardiac mispatterning and AVSD like that observed in *Smo^K399A/K399A^* embryos ^4,53^.

Lung morphogenesis is tightly linked to cardiac development and also requires SHH signaling activity at distinct developmental stages. SHH stimulates initial lung budding from the foregut around E9.5 and contributes to branching morphogenesis as prenatal lung patterning proceeds ^3,5^. Strong and rapid induction of SHH signaling is also necessary during the neonatal period when surfactant secretion occurs, and the alveoli begin to mature ^67^. Failure to induce SHH signaling at this stage leads to alveolar simplification akin to what is observed in *Smo^K399A/K399A^* mice. Moreover, lungs of *Smo^K399A/K399A^* mice had reduced SatPC levels at birth compared to controls, suggesting that AA-mediated allosteric enhancement of SMO signaling may contribute to surfactant secretion or production. Although SHH has not been previously linked to regulation of lung surfactant, AA has been shown to stimulate surfactant release from AT2 cells ^59,60^. Future studies will be required to determine whether these effects result from AA activity in the SHH pathway or if signal synergy occurs between SHH and other AA impacted pathways during this crucial stage of alveolar morphogenesis.

Computer docking analysis placed AA in a binding cavity at the top of the 7TM bundle and predicted that K399 in EC2 of murine SMO anchors AA into this pocket. The prediction suggests that AA, cyclopamine, and the small molecule SMO agonist SAG likely share an overlapping binding site, which is consistent with our published work showing that AA competes with cyclopamine for SMO binding and fails to enhance signaling induced by SAG ^15^. Because the predicted AA pocket is offset from the cholesterol binding cavity in the 7TM domain, we hypothesized that AA and 7TM/IC binding sterols may occupy their respective pockets at the same time and that AA may function as a “cap” to stabilize sterol binding deep in the 7TM core. Accordingly, AA supplementation enhanced signaling by both CHO:MβCD and the TM/IC loop binding oxysterol 24(S),25-EpCHO while depletion of AA by inhibition of cPLA2α reduced pathway induction by sterols or SHH. AA-mediated enhancement and GIRI-mediated suppression of SMO activation by 7β,27-DHC, which binds to the extracellular CRD, was comparatively modest. Thus, AA binding may not influence SMO responsiveness to CRD-binding agonists to the same extent as it does for sterols and oxysterols that bind the IC loops or deep in the 7TM pocket. Notably, the AA anchoring residue, which corresponds to K395 of human SMO, is suggested to function as a coordinating residue for “super stabilizer” ligands that place TM helix 4 in a position that promotes communication between the CRD and EC loops ^21,68^. Accordingly, introduction of K399A into the D477R background, which is thought to be constitutively active due to the arginine substitution “locking” cholesterol into the SMO 7TM pocket, failed to attenuate SMO^D477R^ signaling activity ^22^. As such, we propose that AA allostery acts by enhancing sterol ligand retention in the SMO 7TM core and suggest that this mechanism may be active in signaling contexts where rapid and robust SHH pathway activity is required.

## Methods

### Cell culture

Unless otherwise indicated, all cell lines were grown in Dulbecco’s Modified Eagle Medium (DMEM) supplemented with 10% bovine calf serum, 2 mM L-glutamine, 1 mM sodium pyruvate, 1X non-essential amino acids, and 1% penicillin-streptomycin solution (Thermo Fisher Scientific). Cells were routinely passaged using a 0.25% Trypsin/EDTA solution and maintained in a humidified incubator at 37°C with 5% CO_2_. All cell lines were regularly tested for mycoplasma contamination using MycoAlert (Lonza). To promote ciliogenesis, cell lines were serum starved for 2 h in serum-free media (DMEM supplemented with 0.1 mM nonessential amino acids, 2 mM L-glutamine, 1 mM sodium pyruvate, and 1% penicillin-streptomycin solution). After starvation, media was replaced with low serum (0.5%) media and cells were incubated for 18 h.

### Cloning

Murine *Smo* was amplified from pCS2^+^ *Smo-YFP* and inserted in frame to generate the pEF5/FRT-*Smo-YFP* backbone using the using the pEF5/FRT/V5 Directional TOPO™ Cloning Kit (Thermo Fisher Scientific). Site-directed mutagenesis with the QuikChange XL kit (Agilent Technologies) was used to introduce K399A and E522A substitutions into the constructs. K399A, D477R, D477R+K399A, M2, and M2+K399A substitutions were created in the pEF5/FRT-*Smo-YFP* backbone through the custom cloning services of GenScript Biotech.

### Cell line generation

NIH-3T3 *Smo^−/−^* Flp-In cells have been previously described ^69^. Clonal cell lines stably expressing the proteins of interest were developed using the Flp-In system (Thermo Fisher Scientific). pEF5 FRT *Smo*^WT^*-*YFP, *Smo-*YFP*^K399A^*, *Smo^E522A^-*YFP, *Smo^D477R^-*YFP*, Smo ^M2^-*YFP*, Smo^D477R+K399A^ -*YFP, *or Smo ^M2+K399A^ -*YFP and pOG44 Flp-recombinase vector were transfected using Lipofectamine 3000 (Thermo Fisher Scientific) and selected using Hygromycin B. After chemical selection, cell lines were sorted on the BD FACSDiscover™ S8 for mid-low YFP expression. All cell lines were routinely validated using functional assays and western blotting as appropriate.

### Chemicals

Oxysterols (Avanti Polar Lipids) were dissolved in DMSO. Cyclopamine, SAG, and arachidonic acid (Cayman Chemical) were dissolved in ethanol. Giripladib (GIRI) was synthesized as described ^29^. Resuspended chemicals were stored at −20°C.

To prepare water-soluble cholesterol-MCD complexes, 40 mM cholesterol and MβCD stock solutions were made. Cholesterol was dissolved in ethanol and MβCD was dissolved in low serum media. A glass vial was prepared with 2.5 mM of cholesterol from the organic stock solution. Nitrogen gas was streamed over the cholesterol solution until the organic solvent was evaporated completely, generating a thin film in the vial. One mL of 25 mM MβCD was added to the dried cholesterol film in the glass vial. The solution was vortexed to dissolve the mixture until it became clear. Solutions were passed through a 0.2 μm filter and stored in glass vials at 4°C. Unless otherwise stated, the cholesterol:MβCD ratio was 1:10 in inclusion complexes.

### Expression vectors and transient transfection

SSTR3-GFP (Addgene #49098) ^70^ and ARL13B-GFP (Addgene #40879) ^71^ expression constructs were transfected into NIH-3T3 cells 24 h prior to serum starvation and drug treatments. Transfection of plasmid DNA for transient protein expression was performed using Lipofectamine 3000 (Thermo Fisher Scientific) per the manufacturer’s instructions.

### Immunofluorescence

Cells were cultured on coverslips (Corning) for 18 h and then pretreated with either vehicle, GIRI or cyclopamine in serum-free media. After 2 h, the media was replaced with low serum media containing vehicle, GIRI, SHH-N conditioned media (100 µL/mL), SAG (100 nM), cyclopamine (10 µM), oxysterols (30 µM), or cholesterol:MβCD, and incubated for approximately 20 h. Post-incubation, the cells were washed with PBS and fixed in 4% paraformaldehyde for 12 min. Cells were then washed three times with a wash buffer (PBS with 0.1% Triton X-100) for 5 min each and incubated in the blocking buffer (PBS with 2% BSA, 0.1% Triton X-100, 1% goat serum) for 60 min at room temperature (22°C).

Primary antibody incubations were carried out overnight at 4°C using the following antibodies and dilutions: anti-SMO (Santa Cruz, 1:500), anti-acetylated-α tubulin (Cell Signaling, 1:1000), anti-IFT88 (Proteintech, 1:500), anti-p-cPLA2 (Cell Signaling, 1:1000), anti-gamma tubulin (Abcam, 1:1000), and/or anti-ARL13B (BiCell Scientific, 1:1000) in blocking buffer. Secondary antibody incubations were performed with AlexaFluor 488, 555, and 647 (1:1000) conjugated secondary antibodies (Thermo Fisher Scientific) and DAPI for 60 min at room temperature.

Following secondary antibody incubation, coverslips were washed with wash buffer three times for 5 min each and mounted using ProLong Glass Antifade Mountant (Thermo Fisher Scientific). Microscopy images were captured using a TCS SP8 STED 3X confocal microscope (Leica). For all immunofluorescence experiments, ≥75 cells were examined over at least two experiments, with representative images shown.

### Quantification of fluorescence intensities

For ciliary SMO/SSTR3/IFT88/ARL13B signal intensity determination, each cilium surface area was traced from the base to the tip using the spline profile function using LASX software. The average fluorescence measurement adjacent to the cilium was subtracted from ciliary fluorescence to correct for background. The intensity values along the cilium were exported and analyzed via GraphPad Prism version 10 (GraphPad Software, La Jolla, CA, USA).

### Luciferase assays

SHH Light II cells, which stably express a GLI-responsive firefly luciferase reporter and a constitutive TK-renilla ^31^, were pretreated with either GIRI or a vehicle for 2 h at 37°C in serum-free media. Media was then replaced with low serum (0.5%) media containing GIRI, SHH-N conditioned media (100 μl/mL), cholesterol:MβCD, oxysterols, SAG (100 nM), or arachidonic acid as indicated and incubated for 36 h. SHH-N-conditioned media was collected from HEK293T cells stably expressing the amino-terminal signaling domain of SHH ^72^. Luciferase activity was measured using the Dual-Luciferase Reporter Assay system (Promega).

### Ligand docking

The published ligand-bound SMO structures were obtained from the Protein Data Bank entries 5L7D (cholesterol-bound), 6D32 (cyclopamine-bound), and 6O3C (structure of active Smoothened bound to SAG21k, cholesterol, and NbSmo8) and subsequently loaded into Maestro (Schrödinger Release 2019-3). To prepare the protein for docking, the protein preparation wizard was used to assign bond orders, add hydrogens, create disulfide bonds, and fill in missing side chains and loops. Default parameters were used for optimization of hydrogen bond assignment (sampling of water orientations and use of pH 7.0). Waters beyond 5 Å of het groups or with less than three hydrogen bonds to non-waters were removed. Restrained energy minimization was then applied using the OPLS3e forcefield ^73^. Prepared protein systems were further checked with Ramachandran plots to ensure there were no steric clashes. To generate receptor grids for docking, the ligand respective of each structure was first selected as the grid-defining ligand (cholesterol or cyclopamine). Default Van der Waals radius scaling parameters were used (scaling factor of 1, partial charge cutoff of 0.25). Chemical structures of cholesterol, arachidonic acid, and cyclopamine were obtained from the PubChem database (https://pubchem.ncbi.nlm.nih.gov/) as SMILES identifiers, and subsequently loaded into the LigPrep panel of Maestro (to convert to a 3D structure at target pH of 7.0 ± 2.0 and retaining specified stereochemical properties). Ligands were then docked using the most stringent docking mode (extra precision, “XP”) of Glide ^74^, with the following parameters: dock flexibly, perform post-docking minimization, and keep 100% of scoring compounds.

### Sequence alignment

The Clustal W2 program was used to align *Xenopus* (UniProt A0A974HT42), human (UniProt Q99835), mouse (UniProt P56726), rat (UniProt P97698), chicken (UniProt O42224), zebrafish (UniProt Q5RH73) and fruit fly (UniProt P91682) SMO sequences.

### Immunoblotting

Cells were serum-starved for 2 h, followed by overnight treatment with SHH-N in low serum media. Following incubation, cells were then washed once with cold PBS and lysed in RIPA buffer (150 mM sodium chloride, 1.0% NP-40, 0.5% sodium deoxycholate, 0.1% sodium dodecyl sulfate, 50 mM Tris, pH 8.0) supplemented with EDTA-free complete protease inhibitor cocktail tablets (Roche). Lysates were then sheared 10 times with a 23G needle and kept on ice for 30 min at 4°C. Centrifugation at 14,000 x g at 4°C for 20 min was performed post incubation and the supernatant was collected and quantified for protein concentration using the Pierce BCA Protein Assay Kit (Thermo Fisher).

Samples were diluted in 5X SDS sample buffer and were run on 4%–15% Tris-glycine SDS-PAGE gels (Bio-Rad) and transferred onto Immobilon-P PVDF membranes (Millipore) using Tris/Glycine buffer (Bio-Rad) at 100 V for 1 h. Membranes were blocked with 5% milk in Tris-buffered saline with 0.1% Tween-20 (TBST) for 1 h at room temperature (22°C). Primary antibody incubations were performed at 4°C overnight with the following dilutions: mouse anti-SMO (Santa Cruz, 1:500), rabbit anti-Kinesin (KIF5B, Abcam, 1:5000), mouse anti-GLI1 (Cell Signaling, 1:1000), mouse anti-β-actin (Santa Cruz, 1:500), or rabbit anti-GFP (Rockland, 1:4000). Blots were washed three times in TBST buffer and incubated for 1 h with corresponding HRP-conjugated secondary antibodies (Jackson ImmunoResearch Labs, 1:5000). Blots were developed using the Odyssey Fc imaging system (Li-COR) with the ECL Prime chemiluminescent substrate (Cytiva).

### Membrane thermal shift assay (MTSA)

Membranes were prepared from HEK293T cells transfected with *Smo^WT^-*YFP, *Smo ^K399A^-*YFP, and *Smo^E522A^-*YFP using Lipofectamine 3000. Approximately 36 hours after transfection, media was aspirated, and cells were washed with 1X PBS. Cells were collected in hypotonic (HK) lysis buffer (20 mM HEPES, 10 mM KCl, pH 7.9) with PIC (Roche, 1X) and DTT (5 µM), and incubated on ice for 20 min. Cells were then Dounce homogenized (type B pestle) and cleared of nuclei (centrifuged 2,000 *× g* for 15 min at 4°C). Supernatants were centrifuged at 100,000 *× g* for 30 min at 4°C. The resulting supernatant was separated from the membrane enriched pellet. The membrane enriched pellet was resuspended in HK containing 1% NP-40 buffer.

Membranes (5 µg protein/20 µL final reaction volume) were incubated at 37°C with vehicle, cyclopamine (10 µM), or arachidonic acid (12.5 µM) for 1 h and then incubated for 3 min at various temperatures (34°C–64°C) to establish a thermal denaturation curve. Samples were then treated with ice-cold HK buffer supplemented with NP-40 to a final concentration of 0.8% and snap-frozen in liquid nitrogen. Samples were then thawed at 25°C in a thermo shaker before being transferred to ice. The freeze-thaw cycle was repeated, and ultracentrifugation at 100,000 x g was performed for 20 min at 4°C to precipitate the denatured protein. The supernatant was subjected to immunoblotting. For MTSA plots, western blot intensities were obtained by quantifying the chemiluminescence count per square millimeter (I = counts per mm^2^) using ImageJ-win 64 (Fiji) and normalized to intensity at the lowest temperature. GLI1 and SMO intensities were analyzed by densitometry using Fiji/ImageJ and calculated as GLI1:SMO divided by densitometry of β-actin. All experiments were carried out three times except for SMO^E522A^-YFP plus AA, which was evaluated twice. Normalized data were pooled and plotted using GraphPad Prism v10.

### Cell surface biotinylation

*Smo^−/−^* cells stably expressing various *Smo* constructs were seeded at a density of ∼8×10^5^ cells/60-mm dish and allowed to grow for 24 h. Cells were then incubated overnight in culture media supplemented with 0.5% BCS. Forty-eight hours post seeding, cells were washed twice with cold 1x PBS pH 7.4 and incubated for 30 min at room temperature (22°C) in 1 mL of 1x PBS containing 0.5 mg/ml EZ-Link Sulfo-NHS-Biotin (Pierce). Biotinylation was quenched by washing cells twice with cold 1x PBS containing 100 mM Glycine. Cells were harvested and lysed in RIPA buffer. Lysates were incubated with 40 μL of Streptavidin agarose beads (Thermo Fisher Scientific) for 2 h at 4°C. Beads were washed three times with RIPA buffer and bead purified proteins were extracted in 2X sample buffer containing 2 mM free biotin. Proteins from the supernatant and Streptavidin-purified beads were analyzed by SDS-PAGE and immunoblotting.

### Deglycosylation and phosphatase treatments

RIPA lysates from *Smo^−/−^* Flp-In cells were incubated with 1000 U of peptide-N-glycosidase F (PNGase F) or 1000 U of endoglycosidase H (EndoH) for 2 h at room temperature before being subjected to SDS-PAGE and western blot analysis. All enzymes were sourced from New England Biolabs.

### Mouse models

Data and materials generated from animals were obtained in accordance with the IACUC-approved St. Jude Children’s Research Hospital protocol 608-100616-10/19. All animal husbandry and procedures were performed in accordance with protocols approved by St. Jude Children’s Research Hospital (SJCRH).

*cPLA2α^−/−^* mice have been previously described ^43^.

### Smo^K399A^ mouse generation

The *Smo^K399A^* mouse model was created using CRISPR-Cas9 technology and direct zygote injection at the SJCRH Genetically Engineered Mouse Model (GEMM) Core Unit. Briefly, a mixture of the 40 ng/µL 3X NLS SpCas9 protein (SJCRH Protein Production Core), 20 ng/µL chemically modified sgRNA (CAGE1103.Smo.g2 – 5’-UUGUAGGCUACAAGAACUAU-3’; Synthego), and 10 ng/µL donor ssODN(CAGE1103.g2.sense.ssODN 5’ agatggagactccgtgagtggcatctgttttgtaggctacGCCaactatcggtaccgtgctggctttgtcctggccccaattggcc 3’; IDT AltR modification) was injected into fertilized zygotes as previously described ^75^. Animals were genotyped by targeted deep sequencing using CAGE1103.Smo.F – 5’ ctacacgacgctcttccgatctCACCTGCTCACGTGGTCACTCCCCT 3’ and CAGE1103.Smo.R – 5’ cagacgtgtgctcttccgatctAGAGGAAGGCAGTGGAGCTGGAAGC 3’ at the Center for Advanced Genome Engineering (SJCRH); the resulting sequences were analyzed using CRIS.py as described ^76^. Animals positive for the desired K399A modification were backcrossed to C57BL/6J mice and then bred to homozygosity.

### Mouse immunohistochemistry

Wild-type C57BL/6J (JAX#000664) and *Smo^K399A^* mutant embryos on a C57BL/6 background were collected and prepared for immunohistochemistry at E9.5. Pregnant dams were euthanized, uterine horns were removed, and embryos were dissected in 1X PBS before being rinsed three times. Whole embryos were imaged using a Stereo Microscope (Leica) and somites were counted. The embryos were then fixed in 4% PFA at room temperature (22°C) for 1 h, rinsed three times with 1X PBS, and cryo-protected by moving to 30% sucrose. The next day, embryos were frozen in O.C.T. Compound (Tissue-Tek) on dry ice. Transverse sections of the embryos were cut at a thickness of 10 µm using a Leica Microm CM1950 cryostat. The sections were briefly dried, washed in 1X TBST, and blocked with a buffer containing 2% BSA, 1% goat serum, and 0.1% Triton-X-100 in 1X PBS. Antibodies diluted in the blocking buffer were applied to the sections and incubated overnight at room temperature in a humidified chamber. The following antibodies and their dilutions were used: mouse anti-PAX6 (1:10, DSHB, PAX6), rabbit anti-OLIG2 (1:300, Millipore, AB9610), rabbit anti-FOXA2 (1:250, Abcam, ab108422), and rabbit anti-NKX2.2 (1:200, Novus Bio, NBP1-82554). After removing the primary antibodies, the sections were washed three times with 1X TBST, then incubated with secondary antibodies (Invitrogen) at a 1:500 dilution for 3 h. The sections were washed three times with 1X TBST, rinsed with tap water, dried, and mounted with cover slips using ProLong Gold mounting media. Imaging was performed using a Leica DMi8 widefield microscope and processed with LAS X software. A minimum of five sections each from at least three embryos per genotype were analyzed. Embryos of matching somite numbers were compared to each other. Analyses were performed on pairs of embryos ranging in number from 25-29.

All images analyzed for neural tube patterning were from the cardiac region adjacent to somites 2 and 3. The branchial pouches, aortic sac, and the size and morphology of the atria and ventricles were used as landmarks to identify and align the transverse sections at the cardiac level. Reference images g and h from the Kaufman Atlas of Mouse Development Plate 19b illustrate the cardiac level regions analyzed for expression domain quantifications ^77^.

### Neural tube progenitor domain quantification

Using the Segmented Line Tool in ImageJ, both the neural tube area and the indicated progenitor domain areas were determined. The expression domain areas were then normalized to the total neural tube area for each section analyzed ^29^. Analysis was performed on three to five embryos per genotype, with 4–6 sections examined per embryo.

### RNA sequencing

Mature *Smo^K399A/+^* females were crossed with *Smo^K399A/+^* males to generate litters of E9.5 *Smo^+/+^* and *Smo^K399A/K399A^* embryos. Multiple litters were harvested and pairs of littermates were used to maximize genetic variability (*n*=4 embryos per genotype analyzed). Pregnant dams were harvested, embryos were dissected, and each embryo was placed in a 1.5-mL Eppendorf tube and flash frozen in liquid nitrogen, then stored at −80°C while genotyping was performed. Once four pairs of embryos were generated, all samples were purified at the same time. Each whole embryo was homogenized, and RNA was extracted by using a RNeasy Micro Kit (Qiagen) following the manufacturer’s protocol. RNA sequencing libraries for each sample were prepared with 1 mg total RNA by using the Illumina TruSeq RNA Sample Prep v2 Kit per the manufacturer’s instructions, and sequencing was completed on the Illumina NovaSeq 6000. The 100-bp paired-end reads were trimmed, filtered for quality (Phred-like Q20 or greater) and length (50-bp or longer), and aligned to a mouse reference sequence GRCm38 (UCSC mm10) by using CLC Genomics Workbench v12.0.1 (Qiagen). The transcript per million (TPM) counts were generated from the CLC RNA-Seq Analysis tool. The differential gene expression analysis was performed by using a non-parametric analysis of variance (ANOVA) using the Kruskal-Wallis and Dunn’s tests on log-transformed TPM between four biological replicates from each of two experimental groups, implemented in Partek Genomics Suite v7.0 software (Partek Inc.). The gene sets enrichment and pathway analyses were performed using GSEA ^78^.

### Quantitative reverse transcriptase polymerase chain reaction (qRT-PCR)

Total RNA was extracted from cells using the TRIzol™ (Invitrogen) method according to manufacturer’s instruction. One thousand nanograms of RNA was used to synthesize complementary DNA (cDNA) using High-Capacity cDNA Reverse Transcription Kit (Applied Biosystems). qRT-PCR reactions were performed on a QuantStudio 7 Flex PCR machine using PowerUp Sybr Green Master Mix (Applied Biosystems). Corresponding changes in the expression levels of *Gli1* was calculated using the ΔΔC_T_ method relative to housekeeping genes, *Ppia*, and *Btf3* ^15^.

### PDGFRA^+^ cell isolation from embryonic lungs

Lungs from E18.5 *Smo^K399A/K399A^* and *Smo^+/+^* littermates were harvested, and lung tissue was placed in pre-warmed DMEM with collagenase IV (310 U/mL). Lungs were minced using scissors and incubated at 37°C for 1 h. Following digestion, an equal volume of DMEM supplemented with 2% FBS was added, and the samples were filtered through a 70-µm cell strainer. The resulting cell suspension was centrifuged at 200 x g for 5 min at 4°C, and the pellet was treated with RBC lysis buffer (Invitrogen) for 5 mins at room temperature. After a second centrifugation, cells were resuspended in purification buffer (PBS with 0.2% BSA and 2% FBS) at a concentration of 1×10^6^/ml. The cell suspension was incubated with biotin-conjugated PDGFRA antibody (1:100, R&D Systems) at room temperature for 10 mins, followed by the addition of streptavidin magnetic beads (Pierce). This cell suspension was then incubated at 4°C for 1 h on a rotator. After incubation, the streptavidin beads were washed three times for 5 min each with wash buffer (PBS with 2% FBS) at room temperature (22°C). TRIzol™ was added directly to the beads and RNA was extracted according to the manufacturer’s instructions. PDGFRA-positive cell enrichment was validated by analyzing *Pdgfra* expression levels in RNA isolated from pre-conjugated cell suspension compared to post-purification. *Gli1* expression levels were analyzed in enriched PDGFRA*+* positive lungs cells.

### Embryo histology for cardiac phenotyping

*Smo^+/+^, Smo^K399A/+^*, and *Smo^K399A/K399A^* E14.5 embryos were dissected and fixed in 4% paraformaldehyde overnight at 4°C. Embryo trunk tissues were then dehydrated, embedded in paraffin, and sectioned at a thickness of 5 µm prior to hematoxylin and eosin (H&E) staining by The Human Tissue Resource Center at The University of Chicago facility. A genotype-blind approach was used for cardiac phenotyping and images were taken using a Leica DM2500 optical microscope.

To analyze lung morphology in adult mice, lungs were dissected and embedded in paraffin blocks, sectioned, and stained with H&E for histology imaging. The vacuole area within randomly chosen lung sections was calculated using Fiji/ImageJ. A threshold on images was set to segment the void spaces and the % area was calculated. Two randomly chosen areas from each lung section were analyzed. Sections from five mice/genotype were included in analysis.

### Lung function measurement

#### Assessment of airway hyperresponsiveness by whole body plethysmography (WBP)

Unanesthetized mice were randomly placed into an 8-chamber plethysmograph and allowed to acclimate for 30 minutes before collecting measurements. To evaluate the change in airway physiological parameters, the pressure within each of the plethysmograph chambers was compared to a reference chamber. Following baseline measurements, breathing frequency, tidal volume, inspiratory, and expiratory events were simultaneously recorded in each chamber at intervals of five minutes for a duration of 45 minutes. The enhanced pause (Penh) parameter was calculated using Buxco Finepointe software (Data Sciences International).

#### Assessment of lung resistance and dynamic lung compliance in response to Methacholine inhalation

Lung resistance (RI) and dynamic lung compliance to increasing doses of nebulized methacholine were measured using the Buxco Finepointe Resistance and Compliance system (Data Sciences International). Anesthetized mice were mechanically ventilated through endotracheal intubation. Gradient doses of methacholine (MeCH: 3.125, 6.25, 12.5, and 25 mg/mL) were nebulized to the heated chamber. Airway resistance and dynamic lung compliance were measured over 3 min with a 1 min recovery period at each MeCH dose. Results were collected and quantified using Buxco Finepointe software (Data Sciences International).

### Saturated phosphatidylcholine (SatPC) analysis

Lung tissue was collected from fetal (E18.5-19.5) and newborn (P0) mice and flash frozen. Lipids were extracted from homogenized lung tissue by the Bligh and Dyer method ^79^. SatPC was isolated using the osmium tetroxide-based method of Mason et al. ^80^ and quantitated by phosphorous measurement, as we have previously described ^81^. SatPC levels reported here represent the total amount of lung SatPC in the lung normalized to lung weight.

### Magnetic resonance imaging

Magnetic resonance imaging (MRI) was performed with a Bruker Clinscan 7T MRI system (Bruker Biospin MRI GmbH). Prior to imaging, mice were anesthetized in a chamber with 3% Isoflurane in oxygen at a flow rate of 1 L/min. Mice were maintained in an anesthetized state with 1%–2% Isoflurane via nose-cone delivery system with a flow rate of 1 L/min. Thermal support was provided to the animals using a heated bed with warm water circulation, and a physiological monitoring system tracked their breath rate. During the MRI procedure, a mouse brain volume coil was positioned over the mouse head, with the mouse placed inside a 72-mm transmit/receive coil.

The imaging protocol included T2-weighted turbo spin echo sequences in horizontal (TR/TE = 3080/40 ms, matrix size = 256 x 256, field of view = 36 mm x 36 mm, slice thickness = 0.5 mm, number of slices = 20), coronal (TR/TE = 4760/42 ms, matrix size = 192 x 192, field of view = 15 mm x 15 mm, slice thickness = 0.5 mm, number of slices = 30), and sagittal (TR/TE = 3580/39 ms, matrix size = 128 x 256, field of view = 16 mm x 32 mm, slice thickness = 0.5 mm, number of slices = 24) orientations.

MRI image analysis was performed using 3D Slicer (Surgical Planning Laboratory). Coronal images were segmented by manually outlining regions of interest to delineate the entire brain. This segmentation process was repeated across all slices to cover the entire brain volume. The brain volume was then calculated based on voxel dimensions. Similarly, a separate segmentation was created for the cerebellum, and its volume was determined using the same method.

### Computed tomography

Computed tomography (CT) scans were conducted using a Siemens Inveon PET/CT system at an isotropic resolution of 45 µm. Mice were anesthetized and positioned as described above. The total duration of the scan was 16 min and 34 sec. The CT images were analyzed using Inveon Research Workplace software (Siemens). Measurements of L1 to L10 dimensions were taken based on predetermined measurement locations.

### Statistical methods

All statistical analyses were performed using GraphPad Prism v10. Student’s t-test was carried out to compare statistical significance between two means, and one-way unpaired non-parametric ANOVA was performed for comparison of multiple means. Overall Chi-square analysis was performed to determine significance between expected and observed Mendelian ratios between different genotypes and correlation of heart defects to the mouse genotypes.

Statistics for ciliary quantification were calculated from at least two independent experiments with at least 50 cilia per condition per experiment. All quantified data are presented as mean ± SD, with p < 0.05 considered statistically significant. Significance depicted as *p < 0.05, **p < 0.01, ***p < 0.001, ****p <0.0001 and ns = not significant.

## Supporting information

Reagent Information

## Materials availability

Plasmids generated in this study will be deposited to Addgene at the time of peer reviewed publication. Cell lines and mouse models will be available upon request to the lead contact pursuant to SJCRH material transfer agreements.

## Data and code availability

The RNAseq dataset was deposited into GEO with an accession number of GSE283994. Raw data will be available upon request to the corresponding author.

## Acknowledgements

We thank members of the Ogden lab, Pao-Tien Chuang (UCSF), Andrew Huber (SJCRH) and Dristin Wiggins (SJCRH) for technical assistance, thoughtful discussion, and experimental advice. D. D’Amore provided editorial assistance on the manuscript. *cPLA2α^−/−^* mice were provided by Dr. Joseph Bonventre (Harvard Stem Cell Institute, Boston, MA). RNAseq experiments were performed by the Hartwell Center at SJCRH. Lung sectioning was performed by the Comparative Pathology Core at SJCRH. Model figures were generated using BioRender. Studies were supported by National Institute of General Medical Sciences grants R35GM122546 (S.K.O.) and by ALSAC of St. Jude Children’s Research Hospital. The Center for Advanced Genome Engineering and Genetically Engineered Mouse Model Shared Resource are supported in part by the National Cancer Institute grant P30CA021765. The content is solely the responsibility of the authors and does not necessarily represent the official views of the National Institutes of Health or other funding agencies.

## Author Contributions

Conceptualization, S.S.A. and S.K.O.; Methodology, S.S.A., W.C.W., M.E.D., M.G., D.P.S., C.A.D., S.P.M., I.P.M., J.P.B., and S.K.O.; Validation, S.S.A., M.E.D., W.C.W.; Analysis, S.S.A., M.G., Y.D.W., J.S., R.C., I.P.M., J.P.B., and W.C.W.; Investigation, S.S.A, M.E.D., D.P.S., R. C., I.P.M., J.P.B, M.G., J.S., Y.Z., and W.C.W.; Resources, S.S.A., M.E.D., D.P.S., P.G.T., S.P.M., J.P.B., I.P.M., and S.K.O.; Writing – Original Draft, S.S.A. and S.K.O.; Writing – Review & Editing, S.S.A., W.C.W., T.C., Y.D.W., P.G.T., I.P.M., J.P.B., M.E.D., C.A.D., and S.K.O.; Visualization, S.S.A.; M.E.D., R.C.; Supervision, S.P.M, P.G.T., T.C., I.P.M., J.P.B., and S.K.O.

## Reagent Requests

Reagents will be available upon request to pursuant to SJCRH material transfer agreement.

## Animal Use

Materials developed from animals were generated per SJCRH/IACUC approved protocol 608-100616-10/19.

