## Supplementary Figures and Legends for "Receptor Allostery Promotes Context-Dependent Sonic Hedgehog Signaling During Embryonic Development"

**Supplementary Figure Legends**

**Supplementary Figure 1. cPLA2α inhibition reduces agonist-induced SMO ciliary enrichment.** (A-B) Confocal images of NIH-3T3 cells show SMO (green) ciliary translocation following treatment with DMSO vehicle or the indicated agonists and vehicle or GIRI. Ac.α-Tubulin (A) and ARL13B (B) mark primary cilia in magenta. DAPI is blue. Scale bar = 10 μm.

**Supplementary Figure 2. SMO-YFP variants harbor complex glycans and reach the cell surface.** (A) Cell lysates were prepared from *Smo*$^{-/-}$ Flp-In-NIH-3T3 cells stably expressing the indicated SMO-YFP variants. Lysates were treated with EndoH, or PNGase as indicated and analyzed by western blot. The open arrow marks the EndoH-sensitive ER fraction and closed arrow marks the post-ER form. β-actin is the loading control. (B) Cell surface biotinylation was performed on *Smo*$^{-/-}$ Flp-In-NIH-3T3 cells stably expressing the indicated SMO-YFP proteins. Biotinylated proteins were collected on streptavidin beads, eluted, and analyzed by western blot. Light and dark exposures of the SMO blot are shown. Kinesin was used as the input control. (C) SHH-stimulated SMO ciliary translocation in the indicated cell lines was evaluated by confocal microscopy. The ciliary axoneme is marked by acetylated α-tubulin (magenta) and SMO is shown in green. DAPI (blue) marks nuclei. Scale bar = 5 μm.

**Supplementary Figure 3. Analysis of *Smo*$^{K399A/K399A}$ mice.** (A) A diagram for generation of CRISPR/Cas9 *Smo*$^{K399A}$ knockin mice shows the sgRNA targeting exon 6 of the *Smo* gene (as per the NCBI transcript ID: NM_176996.5). The sgRNA was co-transfected with the Cas9 protein into C57BL/6J zygotes, as described in the materials and methods section. ssODN donor with alanine was used to create the single point mutation at the 399$^{th}$ residue of the SMO protein by homologous recombination. The genotyping primer binding locations in the *Smo*$^{WT}$ and *Smo*$^{K399A/K399A}$ alleles are depicted using black arrows. Depictions are not to scale. (B) The diagram shows MicroCT linear-based measurement landmarks that were used for analysis of skulls along the sagittal and horizontal planes for the *Smo*$^{K399A/+}$ and *Smo* $^{K399AK399A}$ mice. (B') Linear measurements of indicated landmarks (L1-10) were calculated from *Smo*$^{K399A/+}$ and *Smo* $^{K399AK399A}$ mice (8 mice per genotype). Male and female data were combined, and results are shown as mean ± SD. Statistical significance was calculated using a Student's t-test. (C) One of two "escaper" *Smo*$^{K399A/K399A}$;*cPLA2α*$^{-/-}$ mice that survived to weaning does not show an overt phenotype compared to the heterozygous littermate control. (D) Neural tubes of E9.5/25-29 somite stage *Smo*$^{K399A/+}$ and *Smo*$^{K399A/K399A}$ show normal *FoxA2, Nkx2.2, Olig2*, and *Pax6*-expressing progenitor domains. At least 3 sections each from 3 embryos were analyzed. A representative embryo is shown. (E) MRI scans of *Smo*$^{K399A/+}$ and *Smo* $^{K399AK399A}$ brains from adult mice show similar brain morphology. (E') Quantification of cerebellum volume relative to total brain volume was calculated for *Smo*$^{K399A/+}$ and *Smo* $^{K399AK399A}$ mice from MRI scans of both female and male mice. At least 4 mice of each genotype were analyzed. Statistical significance was calculated using a Student's t-test. Error bars indicate SD.

**Supplementary Figure 4. Gene expression analysis of E9.5/25-29 somite stage *Smo*$^{WT}$ and *Smo*$^{K399A/K399A}$ mutant embryos.** Wiki pathways upregulated (A) and downregulated (A') Gene Set Enrichment Analyses (GSEA) for *Smo*$^{K399A/K399A}$ embryos compared to *Smo*$^{+/+}$ are shown. *n*=4 embryos for each genotype. Representation of the positively correlated enrichment of heart development (B) and negatively regulated (B') genes related to ciliopathies in *Smo*$^{K399A/K399A}$ embryos are shown compared to *Smo*$^{+/+}$ control. The top three Gene Ontology (GO) terms of cellular component, biological process and molecular function upregulated (C) and downregulated (C') in *Smo*$^{K399A/K399A}$ embryos compared to *Smo*$^{+/+}$ are shown. (D) A volcano plot with the log10 of the p-value versus the log ratio fold-change in differentially expressed genes (DEGs) in *Smo*$^{K399A/K399A}$ embryos compared to *Smo*$^{+/+}$ are shown. Genes with p-value < 0.01 and log2ratio > 0.5 (upper right quadrant) and < 0.5 (upper left quadrant) are shown. (E) A heatmap of the top statistically significant differentially regulated genes shows ranking by expression value in log scale. (F-F’) Top three GO terms for statistically significant differentially regulated genes in *Smo*$^{K399A/K399A}$ embryos as compared to *Smo*$^{+/+}$ with p< 0.05 (302 genes) using EnrichR. The x-axis represents the false discovery rate (FDR), which refers to the proportion of false discoveries among a set of hypothesis tests. The circle size indicates the number of DEGs that are associated with each significant pathway. The circle color indicates the significant level with the adjusted p-value <0.05. (G) qRT-PCR analyses showing *Pdgfra* expression in E18.5 PDGFRA$^{+}$ cells isolated from lungs of *Smo*$^{+/+}$ and *Smo*$^{K399A/K399A}$ mutant mice. Elevated *Pdgfrα* expression in the anti- PDGFRA$^{+}$ biotin-enriched lung cells following streptavidin pull down confirms successful PDGFRA$^{+}$ cell isolation. Fold-change in expression was determined using the $2^{-\Delta\Delta Ct}$ method. Average fold change was calculated across five *Smo*$^{+/+}$ and four *Smo*$^{K399A/K399A}$ E18.5 lungs analyzed in triplicate. Significance was determined by one-way ANOVA. A p-value of less than 0.05 was considered statistically significant. Significance is denoted as follows: **<0.01, ****<0.0001 and ns, p > 0.05. Error bars indicate SD.

**Supplementary Table 1:** *Smo*$^{K399A/K399A}$*, Smo*$^{K399A/K399A}$*; cPla2α*$^{+/-}$*, and Smo*$^{K399A/K399A}$;*cPLA2α*$^{-/-}$ mutant mice show enhanced perinatal lethality as indicated by skewed Mendelian ratios between birth and weaning. Overall Chi-square analysis was performed to analyze the significance between expected and observed outcomes. *n*=268 mice.

### Supplementary Figure 1

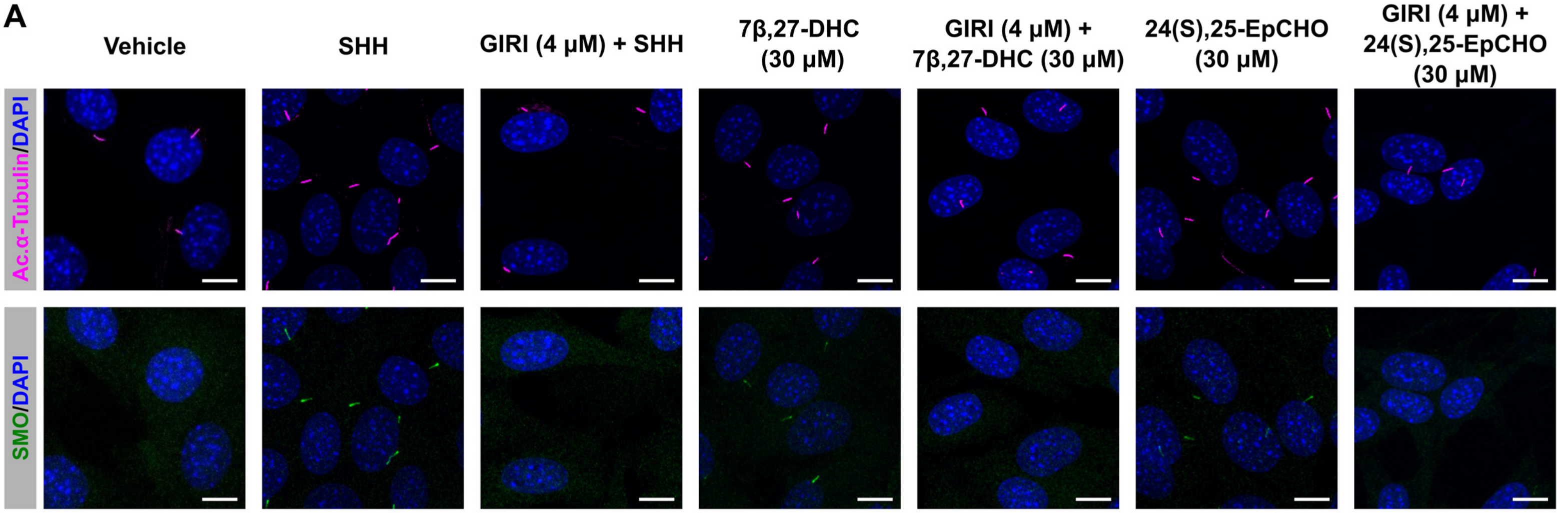


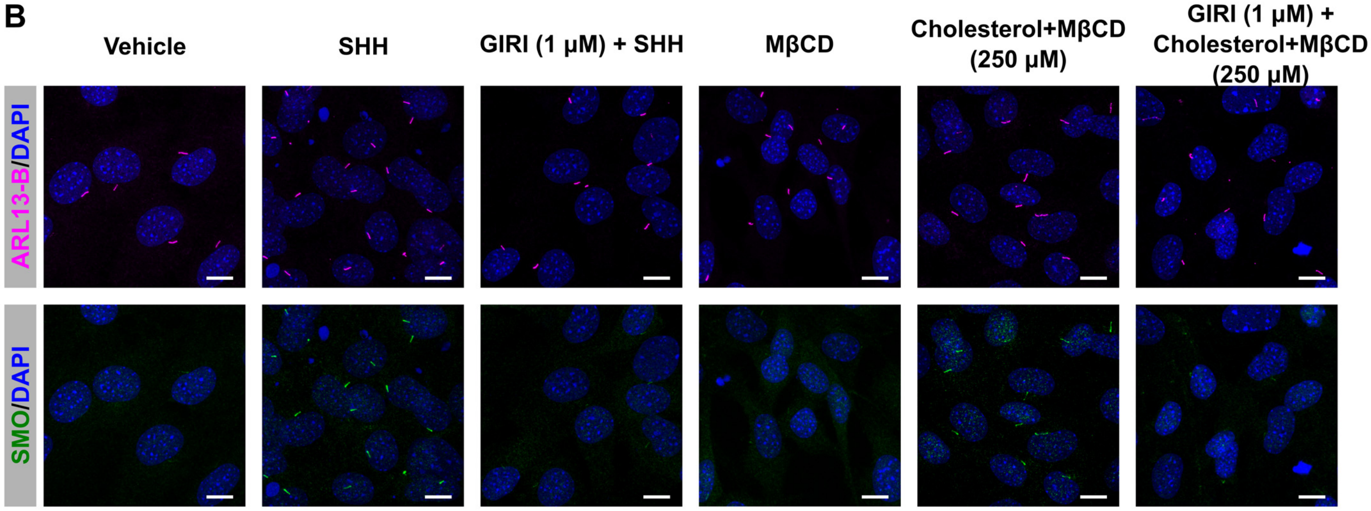

### Supplementary Figure 2

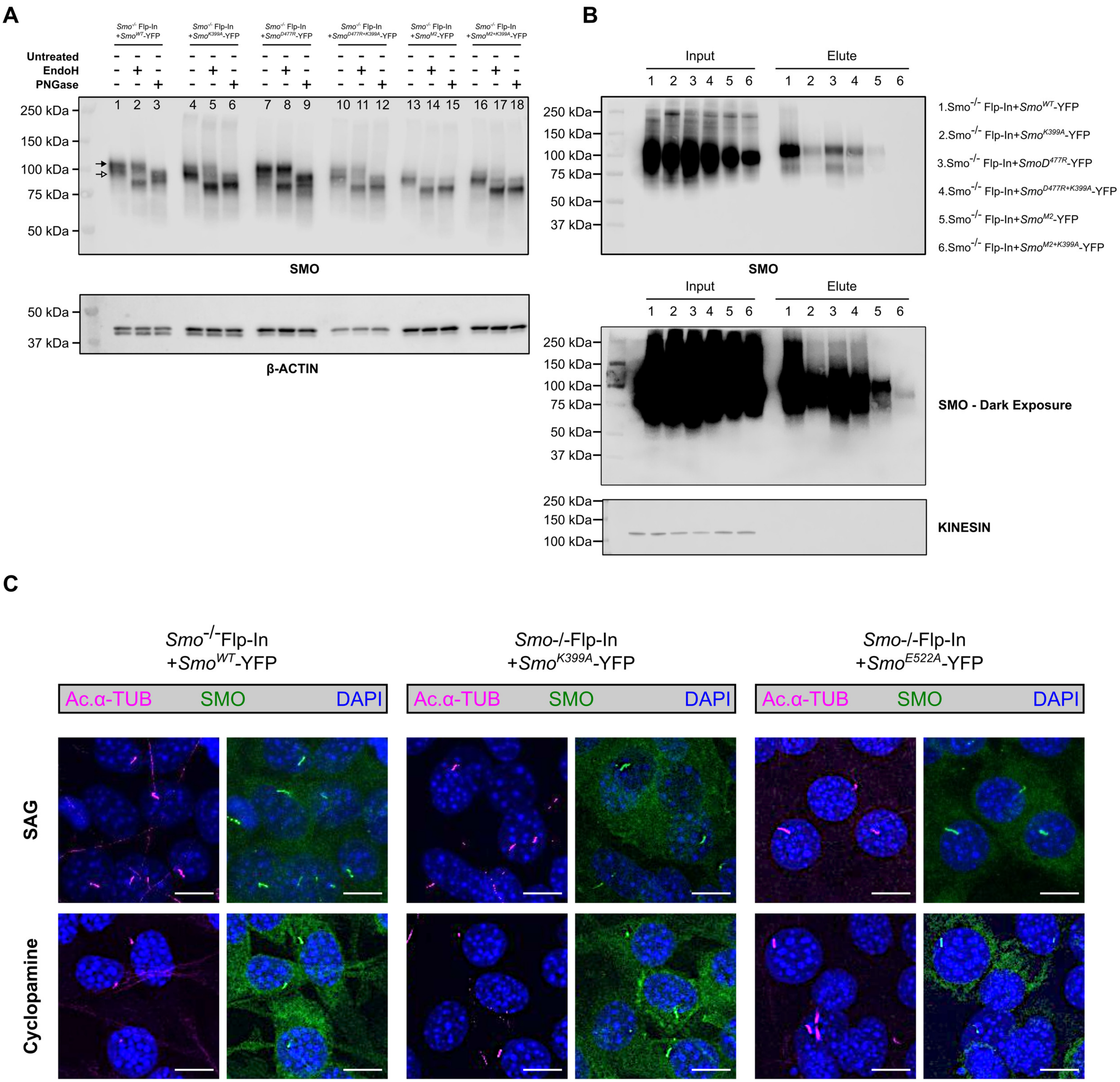

### Supplementary Figure 3

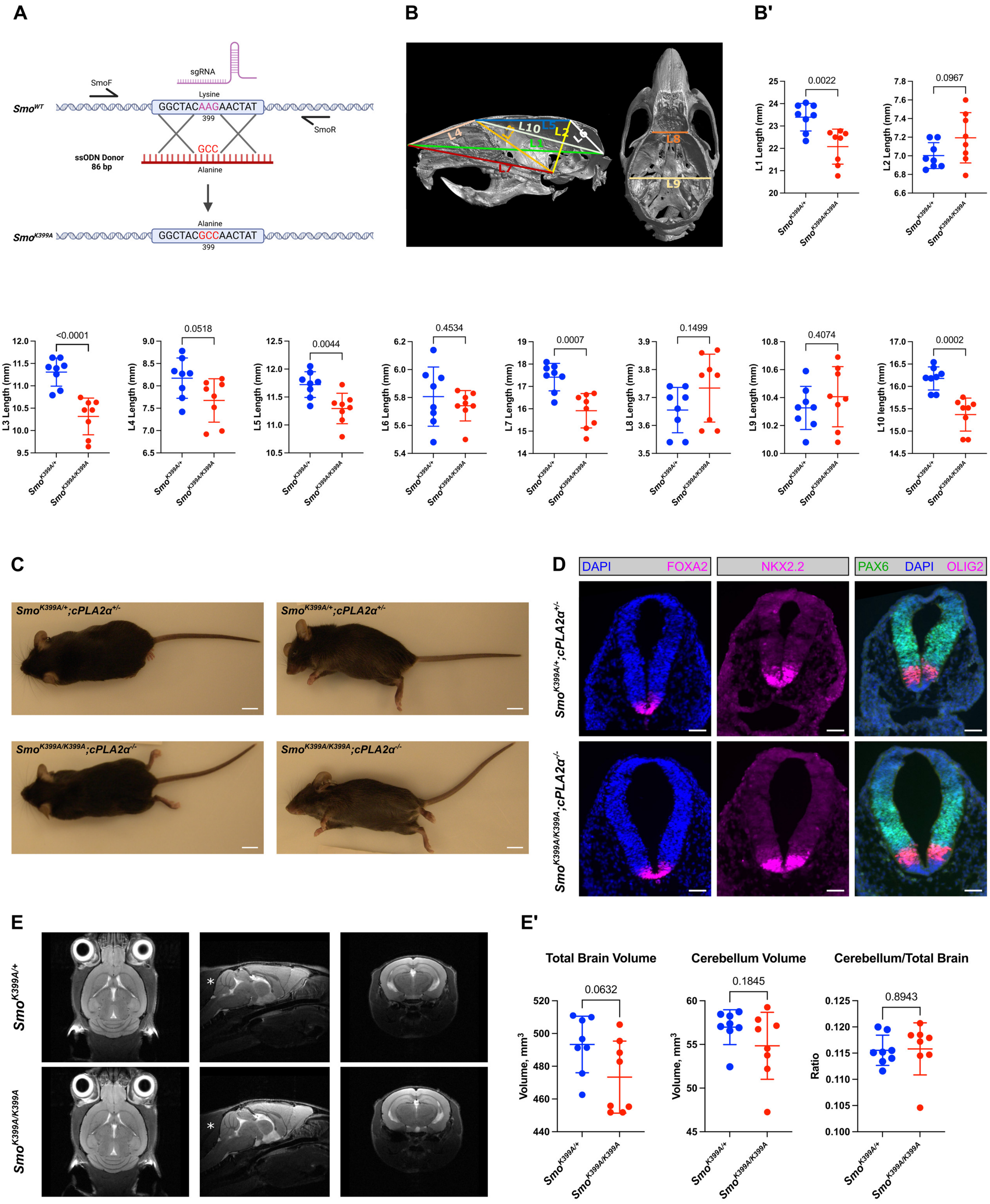

### Supplementary Figure 4

**A**

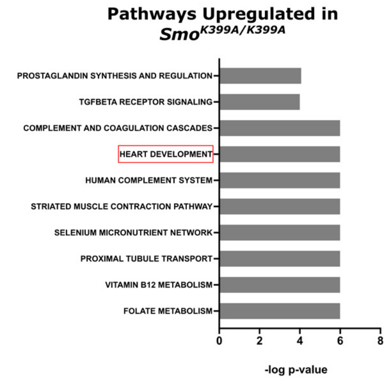


**A'**

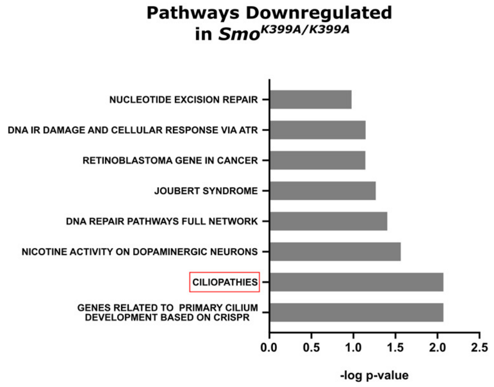


**B**

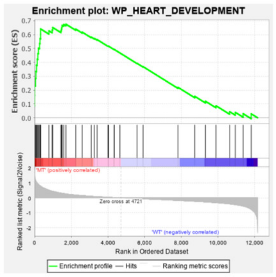


**B'**

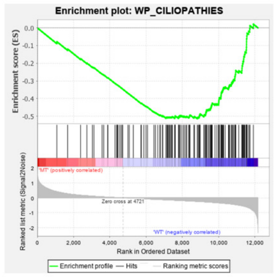


**C**

| | GO TERMS UPREGULATED IN *Smo*$^{K399A/K399A}$ | FDR q- value |
|---|---|---|
| GO Cellular Component | BRUSH BORDER | < 1.0 e-6 |
| | ACTIN FILAMENT BUNDLE | < 1.0 e-6 |
| | CONTRACTILE FIBER | 7.4 e-5 |
| GO Biological Process | PLASMA LIPOPROTEIN PARTICLE CLEARANCE | < 1.0 e-6 |
| | POSITIVE REGULATION OF EPITHELIAL CELL MIGRATION | < 1.0 e-6 |
| | GROWTH FACTOR BINDING | < 1.0 e-6 |
| GO Molecular Function | EXTRACELLULAR MATRIX STRUCTURAL CONSTITUENT | 0.0006 |
| | GROWTH FACTOR BINDING | 0.001 |
| | TRANSFORMING GROWTH FACTOR BETA BINDING | 0.002 |

**C'**

| | GO TERMS DOWNREGULATED IN *Smo*$^{K399A/K399A}$ | FDR q- value |
|---|---|---|
| GO Cellular Component | CENTRIOLE | 0.0006 |
| | REPLICATION FORK | 0.03 |
| | CILIARY BASAL BODY | 0.05 |
| GO Biological Process | CILIARY BASAL BODY PLASMA MEMBRANE DOCKING | 0.007 |
| | AXONEME ASSEMBLY | 0.01 |
| | CILIUM ORGANIZATION | 0.02 |
| GO Molecular Function | CATALYTIC ACTIVITY ACTING ON DNA | 0.05 |
| | DAMAGED DNA BINDING | 0.06 |
| | DEOXYRIBONUCLEASE ACTIVITY | 0.07 |

**D**

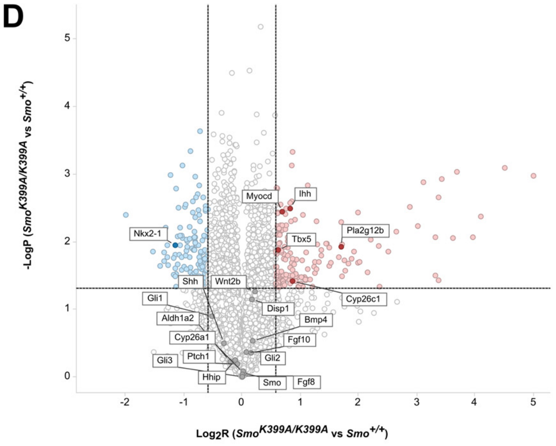


**E**

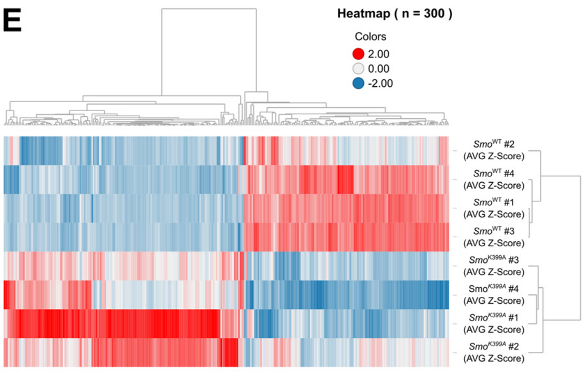


**F**

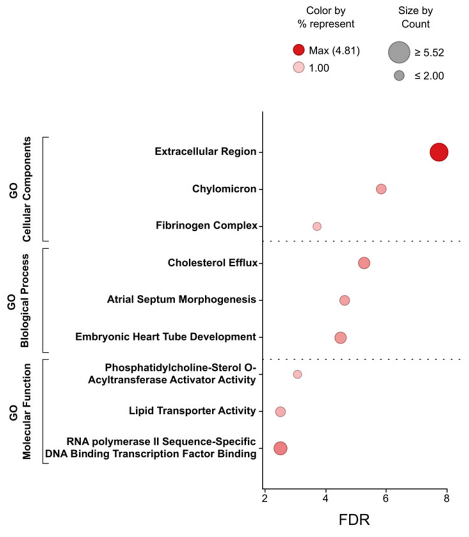


**F'**

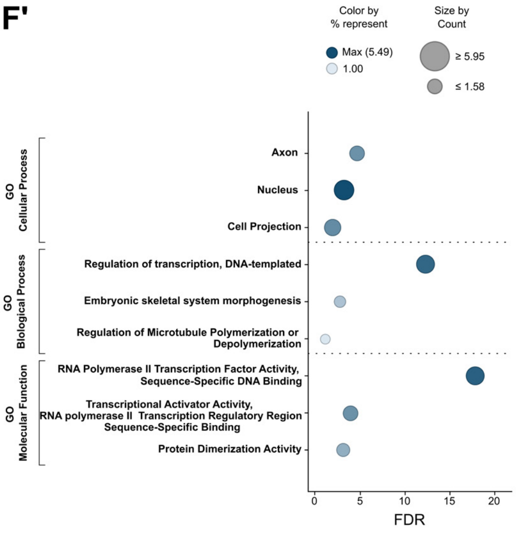


**G**

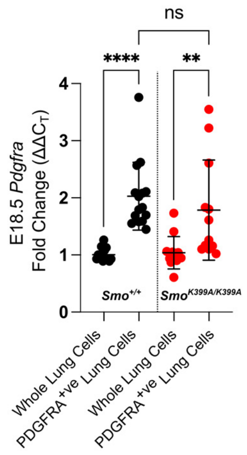

### Supplementary Table 1

## A

| | Expected % | Observed % |
|---|---|---|
| *$Smo^{+/+}$;$cPLA2α^{+/+}$* | 6.25% | 15.67% (42) |
| *$Smo^{+/+}$;$cPLA2α^{+/-}$* | 12.5% | 19.03% (51) |
| *$Smo^{+/+}$;$cPLA2α^{-/-}$* | 6.25% | 6.34% (17) |
| *$Smo^{K399A/+}$;$cPLA2α^{+/+}$* | 12.5% | 18.28% (49) |
| *$Smo^{K399A/+}$;$cPLA2α^{+/-}$* | 25.0% | 25.37% (68) |
| *$Smo^{K399A/+}$;$cPLA2α^{-/-}$* | 12.5% | 10.45% (28) |
| *$Smo^{K399A/K399A}$;$cPLA2α^{+/+}$* | 6.25% | 2.61% (7) |
| *$Smo^{K399A/K399A}$;$cPLA2α^{+/-}$* | 12.5% | 1.49% (4) |
| *$Smo^{K399A/K399A}$;$cPLA2α^{-/-}$* | 6.25% | 0.75% (2) |

**Overall Chi-Square p= 0.005**
