## Supplementary material for "Receptor Allostery Promotes Context-Dependent Sonic Hedgehog Signaling During Embryonic Development": Reagent Information

**REAGENTS**

| REAGENT or RESOURCE | SOURCE | IDENTIFIER |
|---|---|---|
| **Antibodies** | | |
| Mouse monoclonal anti-Smoothened (E-5) | Santa Cruz Biotechnology | Cat# sc-166685; RRID: AB_2239686 |
| Rabbit monoclonal anti-Acetyl-alpha-Tubulin (Ly240) (D20G3) | Cell Signaling Technology | Cat# 5335; RRID: AB_10544694 |
| Rat polyclonal anti-Anti-ARL13B (ARL2L1) | BiCell Scientific | Cat# 90413 |
| Rabbit polyclonal anti-p-cPLA$_2$ (Ser505) | Cell Signaling Technology | Cat# 2831; RRID: AB_2164445 |
| Mouse monoclonal anti-GLI1 (L42B10) | Cell Signaling Technology | Cat# 2643; RRID: AB_2294746 |
| Rabbit monoclonal anti-IFT88 | Proteintech | Cat# 13967-1-AP; RRID: AB_2121979 |
| Mouse polyclonal anti- PDGFR alpha Biotinylated | R&D Systems | Cat# BAF1062; RRID: AB_2162051 |
| Mouse monoclonal anti-PAX6 | DSHB | Cat# pax6; RRID: AB_528427 |
| Rabbit polyclonal anti-OLIG2 | Millipore | Cat# AB9610; RRID: AB_570666 |
| Rabbit polyclonal anti-FOXA2 | Abcam | Cat# ab108422; RRID: AB_11157157 |
| Rabbit polyclonal anti-NKX2.2 | Novus | Cat# NBP1-82554; RRID: AB_11005513 |
| Mouse monoclonal anti-gamma-tubulin | Abcam | Cat# ab11316; RRID: AB_297920 |
| Rabbit monoclonal anti-KIF5B | Abcam | Cat# ab167429; RRID: AB_2715530 |
| Mouse monoclonal anti-beta-Actin (AC-15) | Santa Cruz Biotechnology | Cat# sc-69879; RRID: AB_1119529 |
| Rabbit monoclonal anti-GAPDH | Cell Signaling Technology | Cat# 5174; RRID:AB_10622025 |
| Rabbit polyclonal anti-GFP | Rockland | Cat#600-401-215 |
| Goat anti-Mouse IgG (H+L) Highly Cross-Adsorbed Secondary Antibody, Alexa Fluor™ 488 | Thermo Fisher Scientific | Cat# A-11029; RRID: AB_2534088 |
| Goat anti-Rabbit IgG (H+L) Highly Cross-Adsorbed Secondary Antibody, Alexa Fluor™ 488 | Thermo Fisher Scientific | Cat# A-11034; RRID: AB_2576217 |
| Goat anti-Mouse IgG (H+L) Highly Cross-Adsorbed Secondary Antibody, Alexa Fluor™ 555 | Thermo Fisher Scientific | Cat# A-21424; RRID: AB_141780 |
| Goat anti-Rabbit IgG (H+L) Highly Cross-Adsorbed Secondary Antibody, Alexa Fluor™ 555 | Thermo Fisher Scientific | Cat# A-21429; RRID: AB_2535850 |
| Goat anti-Mouse IgG (H+L) Highly Cross-Adsorbed Secondary Antibody, Alexa Fluor™ 647 | Thermo Fisher Scientific | Cat# A-21236; RRID: AB_2535805 |
| Goat anti-Rabbit IgG (H+L) Highly Cross-Adsorbed Secondary Antibody, Alexa Fluor™ 647 | Thermo Fisher Scientific | Cat# A-21245; RRID: AB_2535813 |
| Peroxidase-AffiniPure Donkey Anti-Mouse IgG (H+L) | Jackson ImmunoResearch Labs | Cat# 715-035-151; RRID: AB_2340771 |

| | | |
|---|---|---|
| Peroxidase-AffiniPure Donkey Anti-Rabbit IgG (H+L) | Jackson ImmunoResearch Labs | Cat# 711-035-152; RRID: AB_10015282 |
| **Chemicals, peptides, and recombinant proteins** | | |
| Arachidonic Acid | Cayman Chemical | Cat# 90010-50 |
| Giripladib | WuXi Custom Synthesis | DOI: 10.1021/acs.biochem.5b00549 |
| SAG | Selleck Chemical | Cat# S7779-2MG |
| Cholesterol | EMD Millipore | Cat# 228111-5GM |
| 20-Hydroxy Cholesterol | Cayman Chemical | Cat# 20103 |
| 7β,27-dihydroxycholesterol (7β,27-DHC) | Abcam | Cat# AB141633-1MG |
| 24,25-epoxycholesterol (24,25-EC) | Avanti Polar Lipids | Cat# 700025P-1MG |
| Methyl-beta-cyclodextrin | Sigma-Aldrich | Cat# C4555-1G |
| Cyclopamine | LC Laboratories | Cat# C-8700 |
| PNGase F | New England Biolabs | Cat# P0704S |
| Endo H | New England Biolabs | Cat# P0702L |
| DMEM media | Life Technologies | Cat# 11960-044 |
| Bovine Calf Serum | Fisher Scientific | Cat# SH30087.03 |
| L-Glutamine | Life Technologies | Cat# 25030081 |
| Sodium Pyruvate | Life Technologies | Cat# 11360-070 |
| Penicillin-Streptomycin (10,000 U/mL) | Life Technologies | Cat# 15140-122 |
| MEM Non-Essential Amino Acids Solution | Life Technologies | Cat# 11140-050 |
| Hygromycin B (50 mg/mL) | Life Technologies | Cat# 10687-010 |
| Paraformaldehyde | Electron Microscopy Solutions | Cat# 15710 |
| ProLong™ Diamond Antifade Mountant | Thermo Fisher Scientific | Cat# P36961 |
| ProLong™ Glass Antifade Mountant | Thermo Fisher Scientific | Cat# P36980 |
| Normal Goat Serum | Jackson ImmunoResearch Labs | Cat# 005-000-121; RRID: AB_2336990 |
| Dulbecco’s Phosphate-Buffered Saline | Corning | Cat# 20-031-CV |
| DAPI (4',6-Diamidino-2-Phenylindole, Dilactate) | Thermo Fisher Scientific | Cat# D3571 |
| DMSO | Sigma-Aldrich | Cat# D2650 |
| Collagenase Type IV | Thermo Fisher Scientific | Cat# 17104019 |
| 10X RBC Lysis Buffer | Invitrogen | Cat# 00-4300-54 |
| 70 µm cell strainer | Genesee Scientific | Cat# 25-376 |
| Biotin | Sigma-Aldrich | Cat# B4501 |

| | | |
|---|---|---|
| Pierce™ Streptavidin Magnetic Beads | Thermo Fisher Scientific | Cat# 88817 |
| Lipofectamine™ 3000 Transfection Reagent | Thermo Fisher Scientific | Cat# L3000001 |
| TRIzol™ | Invitrogen | Cat# 15596026 |
| EZ-Link Sulfo-NHS-Biotin | Thermo Fisher Scientific | Cat# 21217 |
| 10x Tris/Glycine | BioRad | Cat# 1610771 |
| 10x Tris/Glycine/SDS | BioRad | Cat# 1610772 |
| 4-15% TGX Gels, 26 well | BioRad | Cat# 5671085 |
| 7.5%BioRad TGX Gels, 26 well | BioRad | Cat# 5671025 |
| 4-15% TGX Gels, 18 well | BioRad | Cat# 5671084 |
| 7.5%BioRad TGX Gels, 18 well | BioRad | Cat# 5671024 |
| Immobilon-P PVDF Membrane | EMD Millipore Corp. | Cat# IPVH00010 |
| Glycine | Sigma-Aldrich | Cat# G8898 |
| 10x RIPA Lysis Buffer | Millipore | Cat# 20-188 |
| Triton X-100 | Fisher Scientific | Cat# X100-100ML |
| NP-40 | Sigma-Aldrich | Cat# 74385 |
| Tween-20 | Sigma-Aldrich | Cat# P1379 |
| DTT | Sigma-Aldrich | Cat# 43816 |
| Sucrose | Fisher Scientific | Cat# S53 |
| BSA | Fisher Scientific | Cat# BP9700100 |
| PBS | Fisher Scientific | Cat# 21-031-CV |
| 10x TBS | Fisher Scientific | Cat# AAJ60764K7 |
| **Critical commercial assays** | | |
| BCA Protein Assay Kit | Thermo Scientific™ Pierce™ | Cat# 23227 |
| Flp-In™ Complete System | Thermo Fisher Scientific | Cat# K601001 |
| pEF5/FRT/V5 Directional TOPO® Expression Kit | Thermo Fisher Scientific | Cat# K6035-01 |
| ECL™ Prime Western Blotting System | Millipore Sigma | Cat# GERPN2232 |
| Myco Alert Plus | Lonza | Cat# LT07-710 |
| RNeasy Micro Kit | Qiagen | Cat# 74004 |
| TruSeq RNA Library Prep Kit v2 | Illumina | Cat# RS-122-2001 |
| Quickchange II XL Kit | Agilent | Cat# 200522 |
| PowerUp Sybr Green Master Mix | Applied Biosystems | Cat# A25742 |
| Dual-Luciferase Reporter Assay System | Promega | Cat# E1960 |
| High-Capacity cDNA Reverse Transcriptase Kit | Applied Biosystems | Cat# 4368814 |

| | | |
|---|---|---|
| **Deposited data** | | |
| Bulk *Smo*$^{K399A}$ RNA seq | This paper | GEO: GSE283994 |
| **Experimental models: Cell lines** | | |
| *Homo Sapiens*: HEK293T | ATCC | RRID: CVCL_1926 |
| *Mus musculus*: NIH3T3 | ATCC | RRID: CVCL_0594 |
| *Mus musculus*: Light II | ATCC | RRID: CVCL_2721 |
| *Mus musculus*: NIH3T3 Flp-In | Thermo Fisher Scientific | Cat# R76107 |
| *Mus musculus*: *Smo*$^{-/-}$ NIH3T3 Flp-In | DOI: /10.7554/eLife.61432 | N/A |
| *Mus musculus*: *Smo*-YFP$^{WT}$ NIH3T3 Flp-In | This paper | N/A |
| *Mus musculus*: *Smo*-YFP$^{K399A}$ NIH3T3 Flp-In | This paper | N/A |
| *Mus musculus*: *Smo*-YFP$^{E522A}$ NIH3T3 Flp-In | This paper | N/A |
| *Mus musculus*: *Smo*-YFP$^{M2}$ NIH3T3 Flp-In | This paper | N/A |
| *Mus musculus*: *Smo*-YFP$^{D477R}$ NIH3T3 Flp-In | This paper | N/A |
| *Mus musculus*: *Smo*-YFP$^{M2+K399A}$ NIH3T3 Flp-In | This paper | N/A |
| *Mus musculus*: *Smo*-YFP$^{D477R+K399A}$ NIH3T3 Flp-In | This paper | N/A |
| **Experimental models: Organisms/strains** | | |
| C57BL/6J (Wild type) | JAX | RRID: IMSR_JAX:000664 |
| C57BL/6-cPLA2$\alpha^{+/-}$ | Provided by Dr. Joseph Bonventre DOI: 10.1038/37635 | N/A |
| C57BL/6- *Smo*$^{K399A/K399A}$ | This paper | N/A |
| **Recombinant DNA** | | |
| pcDNA3.1(+) (control vector) | Thermo Fisher Scientific | Cat# V79020 |
| pCDNA3-EGFP | N/A | Provided by Dr. Doug Golenbock RRID: Addgene_13031 |
| pEF5B-FRT-AP-Sstr3-GFP-DEST | DOI: 10.7554/eLife.00 654 | Provided by Dr. Maxence Nachury RRID: Addgene_49098 |
| pcDNA-DEST47-Arl13b-GFP | DOI: 10.1091/mbc.E1 0-12-0994 | Provided by Dr. Tamara Caspary RRID: Addgene_40872 |
| pEF5/FRT/V5-DEST™ Gateway™ Vector | Thermo Fisher Scientific | Cat# V602020 |
| pEF5-FRT-YFP-Smo | This paper | N/A |
| pEF5-FRT-YFP-Smo$^{K399A}$ | This paper | N/A |
| pEF5-FRT-YFP-Smo$^{E522A}$ | This paper | N/A |

| | | |
|---|---|---|
| pEF5-FRT-YFP-Smo$^{M2}$ | This paper | N/A |
| pEF5-FRT-YFP-Smo$^{D477R}$ | This paper | N/A |
| pEF5-FRT-YFP-Smo$^{M2+K399A}$ | This paper | N/A |
| pEF5-FRT-YFP-Smo$^{D477R+K399A}$ | This paper | N/A |
| **Oligonucleotides (5'-3')** | | |
| *Gli1* Forward: CCAAGCCAACTTTATGTCAGGG | Thermo Fisher Scientific | N/A |
| *Gli1* Reverse: AGCCCGCTTCTTTGTTAATTTGA | Thermo Fisher Scientific | N/A |
| *Ppia* Forward: AGCACTGGAGAGAAAGGATT | Thermo Fisher Scientific | N/A |
| *Ppia* Reverse: ATTATGGCGTGTAAAGTCACCA | Thermo Fisher Scientific | N/A |
| *Btf3* Forward: GAACAACATCTCTGGTATTGAAGA | Thermo Fisher Scientific | N/A |
| *Btf3* Reverse: AATGGTGAAGGTGTTTGCTG | Thermo Fisher Scientific | N/A |
| *Pdgfra* Forward: GCAGTTGCCTTACGACTCCAGA | Integrated DNA Technologies | N/A |
| *Pdgfra* Reverse: GGTTTGAGCATCTTCACAGCCAC | Integrated DNA Technologies | N/A |
| *QuickChange Smo K399A primer Forward:* AGCACGGTACCGATAGTTCGCGTAGCCTACAAAACA GATG | Thermo Fisher Scientific | N/A |
| *QuickChange Smo K399A primer Reverse:* CATCTGTTTTGTAGGCTACGCGAACTATCGGTACCGT GCT | Thermo Fisher Scientific | N/A |
| *QuickChange Smo E522A primer Forward:* CTCCTGGTGGCAAGATCAATCTATTTGCCATG | Thermo Fisher Scientific | N/A |
| *QuickChange Smo E522A primer Reverse:* GATTGATCTTCGCCACCAGGAGGCTG | Thermo Fisher Scientific | N/A |
| sgRNA SMO: UUGUAGGCUACAAGAACUAU | Synthego | N/A |
| Donor ssODN: agatggagactccgtgagtggcatctgttttgtaggctacGCCaactatcggt accgtgctggctttgtcctggccccaattggcc | Integrated DNA Technologies | N/A |
| NGS Primer *Smo* Forward: ctacacgacgctcttccgatctCACCTGCTCACGTGGTCACTCC CCT | Integrated DNA Technologies | N/A |
| NGS Primer *Smo* Reverse: cagacgtgtgctcttccgatctAGAGGAAGGCAGTGGAGCTGG AAG | Integrated DNA Technologies | N/A |
| **Software and algorithms** | | |
| Fiji (ImageJ) (v2.9.0) | Schindelin et al. RRID:SCR_002285 | http://fiji.sc |

| | | |
|---|---|---|
| CRIS.py (v2.0) | Connelly et al. | https://github.com/patrickc01/CRIS.py |
| Leica Application Suite X (LAS X) (v4.0.0) | Leica RRID:SCR_013673 | https://www.leica-microsystems.com/products/microscope-software/details/product/leica-las-x-ls/ |
| GraphPad Prism (v10.0) | GraphPad RRID:SCR_002798 | http://www.graphpad.com/ |
| Molecular Signatures Database (GSEA) (v2023.1.Mm) | Subramanian et al. RRID:SCR_016863 | http://software.broadinstitute.org/gsea/msigdb/index.jsp |
| CLC Genomics Server (v23.0.4) | Qiagen RRID:SCR_017396 | https://www.qiagenbioinformatics.com/products/clc-genomics-server/ |
| Inkscape | Inkscape RRID:SCR_014479 | https://inkscape.org/ |
| 3D Slicer | 3D Slicer (RRID:SCR_005619) | http://slicer.org/ |
| Inveon Research Workplace (v3.0) | Siemens | N/A |
| Maestro | Maestro (RRID:SCR_016748) | https://www.schrodinger.com/maestro |
| Partek Genomics Suite (v7.0) | Partek Inc. RRID:SCR_011860 | http://www.partek.com/?q=partekgs |
